# CAR-T-drug conjugate against solid tumor

**DOI:** 10.64898/2026.01.02.696502

**Authors:** Shan Xin, Yanzhi Feng, Feifei Zhang, Jiang Yu, Chuanpeng Dong, Bo Tao, Kaiyuan Tang, Qihao Wu, Binfan Chen, Ping Ren, Nipun Verma, Chuan Liu, Kazushi Suzuki, Xinyu Ling, Daniel Park, Harriet Kluger, Jason M. Crawford, Rong Fan, Jiangbing Zhou, Sidi Chen

## Abstract

Chimeric antigen receptor (CAR)-T cell therapy has been clinically successful in hematologic cancers but faces challenges in solid tumors, primarily due to limited tumor infiltration, immunosuppressive tumor microenvironments (TME), and antigen heterogeneity. While combining CAR-T cell therapy with chemotherapy can enhance antitumor activity, this often leads to substantial systemic toxicity. In this study, we introduce CAR-T-drug conjugate (CAR-T-D-C), a new class of dual-functional therapeutics that effectively addresses these obstacles by integrating click chemistry for precise conjugation of cytotoxic agents onto antigen-specific CAR-T cells. By transforming the hostile TME into an ally, this approach facilitates localized delivery of cytotoxic payload directly to the tumor site, enhancing the overall effectiveness of CAR-T therapy in solid tumors. CAR-T-D-Cs with different CAR-T cell binders exhibit robust antitumor activity across diverse solid tumor models, including both human tumor xenografts and syngeneic models. Spatial transcriptomic studies reveals that CAR-T-D-C achieves improved CAR-T tumor infiltration and functional activation within the TME. Compared to conventional CAR-T therapy, CAR-T-D-C markedly enhances immune cell infiltration, augments effector functions, promotes antigen spreading, amplifies systemic immune responses, and improves overall anti-tumor immunity. CAR-T-D-C represents a versatile therapeutic concept that combines the potency of small molecule drugs and the specificity of CAR-T cells as a 2-in-1 immunochemotherapy for treatment of solid tumors.

## Main

Cancer treatment has substantially advanced over recent decades, notably with the development of antibody-drug conjugates (ADCs) as a targeted therapeutic approach for solid tumors^1^. By combining the specificity of monoclonal antibodies with the potent cytotoxicity of chemotherapeutic agents, ADCs achieve targeted drug delivery, reducing systemic toxicity and improving therapeutic efficacy^2–4^. In certain settings, ADCs still encounter major challenges in solid tumors, including heterogeneous target antigen expression, suboptimal tumor penetration, and evasion mechanisms that impede their sustained efficacy^5^. Importantly, a substantial portion of antigens expressed on tumor cell surfaces are unable to undergo endocytosis, thereby restricting the applicability of ADCs in tumors where endocytosis is essential for internalizing the cytotoxic payload^6, 7^.

Since the 2010s, Chimeric antigen receptor (CAR)-T cell therapy has transformed cancer immunotherapy, particularly in hematologic malignancies, by enabling genetically engineered T cells to recognize and eliminate tumor cells expressing specific surface antigens^8–12^. This mechanism allows CAR-T cells to target a broader range of antigens, including those that ADCs cannot effectively engage^9^. Despite this success, translating the success of CAR-T cells to solid tumors has been challenging, due to factors including the immunosuppressive tumor microenvironment (TME), antigen heterogeneity, and inadequate tumor infiltration^13, 14^. These barriers often reduce the efficacy of CAR-T cells in solid tumors and lead to resistance or relapse. The challenges in currently available treatment modalities including ADC and CAR-T necessitate the development of different classes of therapeutics especially for solid tumors.

In this study, we developed a dual-function CAR-T-drug conjugate (CAR-T-D-C), a therapeutic approach that synergizes the specific targeting capabilities of CAR-T cells and the potent cytotoxic effects of an ADC-like drug conjugate. Specifically, we employed click-chemistry-mediated conjugation to link CAR-T cells with chemotherapy agents such as the FDA-approved cytotoxin monomethyl auristatin E (MMAE) as the chemical payload^15, 16^. This approach utilizes a valine-citrulline (val-cit) linker to attach the payload to the cells, allowing selective cleavage by tumor-associated cathepsin B (CTSB) and enabling site-specific activation of MMAE within the TME^17–19^. We hypothesize that this strategy allows CAR-T cells to have a combinatorial effect via not only directly targeting tumor cells, but also acting as a delivery vehicle that mediates localized drug release for elimination of cancer cells in the tumor bed. This new type of therapeutic chimera has the potential against solid tumors.

## Results

### Efficient generation of CAR-T cells targeting solid tumors through LNP-mRNA immunization and single cell BCR sequencing

An increasing number of tumor-associated antigens (TAA) are being identified as potential targets for immunotherapy in solid tumors. Among these, Carbonic anhydrase 9 (CA9/CAIX) has garnered particular attention due to its overexpression across various solid tumor types, with especially high levels observed in renal cancer (Fig. S1a), emphasizing its potential as a solid tumor-associated antigen for targeted therapies. Although traditional ADCs have demonstrated significant success in targeting tumors and eliciting anti-tumor response, the non-internalizing structural characteristics of CA9 limit its applications in ADC-based approaches (Fig. S1b). While CA9 is a promising target based on its expression profiles, a prior anti-CA9 CAR-T phase I/II trial failed to deliver successful outcome either due to lack of efficacy and toxicity^20, 21^. We converted a previously reported antibody sequence (clone G250, US 9605075B2) into a single-chain variable fragments (scFvs) format and generated anti-CA9 G250 CAR-T cells targeting A498 cells. However, our results indicated that these anti-CA9 G250 CAR-T cells were ineffective at killing A498 cells (Fig. S1c).

To strike an optimal efficacy-toxicity balance, it is necessary to first develop a good CAR binder for CA9. We immunized ATX IgG-humanized mice with lipid nanoparticle (LNP)-encapsulated mRNAs encoding full-length human CA9 (Fig 1a), using a procedure described before^22, 23^. The biophysical integrity of the synthesized CA9 LNP-mRNAs was characterized (Fig. S1d). After immunization, blood samples were collected retro-orbitally to evaluate immune response using ELISA (Fig. S1e). The results demonstrated significant CA9-specific antibody binding ability, indicating a robust humoral response to the CA9 LNP-mRNA immunization (Fig. S1e). Next, we performed single-cell B-cell receptor sequencing (scBCR-seq) on memory B cells isolated from the spleen and lymph nodes of CA9-immunized mice (Fig. 1a). Sequencing analysis of 1,407 CA9-specific B cells yielded 627 paired antibody sequences, providing a comprehensive profile of CA9-elicited memory B cell repertoires, revealing clonal expansion patterns, enriched IgG clonotypes, and V/J segment recombination preferences (Fig. S1f, g). This high-resolution mapping offers valuable insights into the antibody repertoire dynamics following antigen-targeted immunization with LNP-mRNA technology.

**Figure 1.**
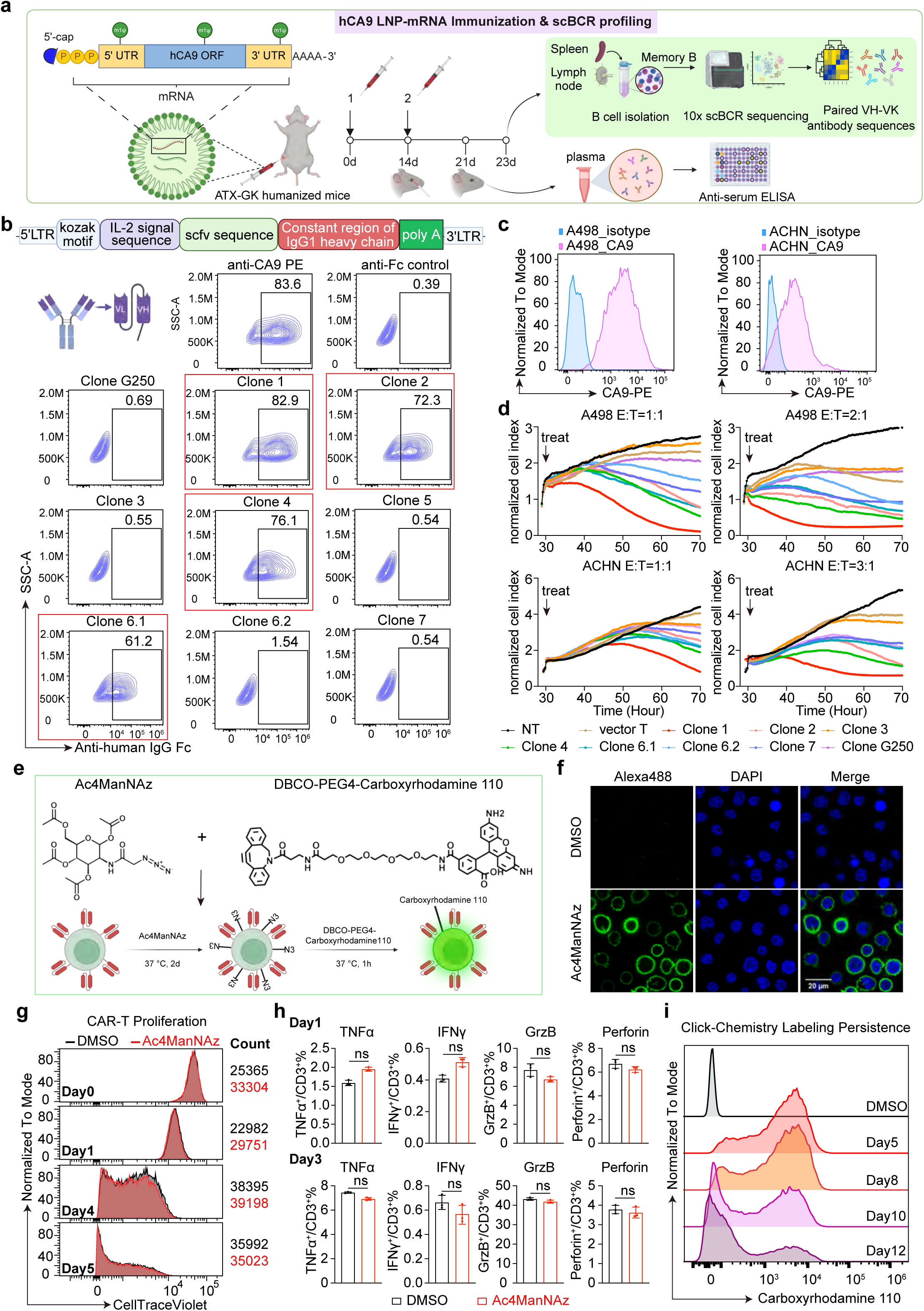
Rapid generation of human anti-CA9 CAR-T binder and efficient click chemistry labeling of anti-CA9 CAR-T cells. **(a)** Schematics of mouse immunization schedule and downstream scBCR profiling. **(b)** The binding ability of the top clones in scFv format was assessed using flow cytometry. A commercial anti-human CA9 antibody (CA9-PE) were used to verify CA9 overexpression. Sequence from a prior binder (clone G250) was used as a positive control. **(c)** Flow cytometry analysis indicating CA9 highly expressed on the surface of A498 and ACHN cell lines. **(d)** Real-Time Cell Analysis (RTCA) killing assays were employed to assess anti-CA9 CAR-T killing ability against A498 **(upper panels)** and ACHN **(lower panels)** cells, as indicated by normalized cell index values. E: T (effector cell to target cell ratio) was indicated. **(e)** Schematics of the metabolic conjugating process of the CAR-T cells with Ac4ManNAz azido-sugar and DBCO-Carboxyrhodamine 110. **(f)** Confocal image demonstrating the successful conjugation of DBCO-PEG4-Carboxyrhodamine 110 on the cell surface of Ac4ManNAz-labeled human CAR-T cells. **(g)** Proliferation of metabolically labeled human CAR-T cells compared to control cells (DMSO), shown using CellTrace Violet labeling. **(h)** Cytokine release of Ac₄ManNAz labeled human CAR-T cells compared with control (DMSO) cells, measured by flow cytometry. **(i)** Histogram showing the fluorescence intensity of Carboxyrhodamine 110-labeled human CAR-T cells overtime. The fluorescence intensity was measured by flow cytometry on days 5, 8, 10 and 12 post-labeling.

To determine the CA9 reactivity of the most enriched BCRs, we tested the top eight pairs of heavy and light variable segments, designated as clones 1-5, 6.1, 6.2, and 7, along with clone G250 as control. Our results showed that 7 of the 8 selected clones, plus the positive control, exhibited strong binding affinity to CA9 in the antibody format, confirming the robustness of our screening approach (Fig. S2a). To assess the compatibility of these clones for CAR-T cell applications, we converted the antibody sequences into scFvs and evaluated their binding to CA9. Among the eight generated scFvs, only four (clone1, clone2, clone4, and clone6.1) retained robust binding to CA9, while the remaining clones, along with the positive control scFv (clone G250), lost binding ability in the scFv format (Fig. 1b).

We cloned all the binding scFvs into CAR constructs containing a 4-1BB costimulatory domain and a FLAG tag. These constructs were then transduced into primary human T cells using adeno-associated virus (AAV) vectors^24^ (Fig. S2b). Flow cytometry analysis confirmed efficient transduction and surface expression of the constructs, with FLAG expression observed in all tested clones (Fig. S2b). FLAG-positive cells were sorted and assessed in a Real-Time Cell Analysis (RTCA) killing assay against CA9-expressing renal cell carcinoma lines A498 and ACHN (Fig. 1c, d). Among the tested CAR constructs, all scFvs showed strong anti-tumor killing ability, with scFv clone 1 showing particularly potent cytotoxicity. Specifically, scFv clone 1 achieved significant killing efficiency against A498 cells at effector-to-target (E: T) ratios of 1:2 and 2:1, and similarly against ACHN cells at E:T ratios of 1:2 and 3:1 (Fig. 1d). Thus, we selected anti-CA9 scFv clone 1 as the lead for downstream studies.

### Precise surface glycolabeling of CAR-T cells via DBCO-azide click chemistry

We next explored the feasibility of directly conjugating drug to our engineered anti-CA9 CAR-T cells through a precision glycolabeling approach, as outlined in the workflow (Fig.1e). Initially, CAR-T cells were incubated with 20 µM Ac4ManNAz for 48 hours to introduce azide groups on the cell surface. After incubation, cells were thoroughly washed and subsequently treated with DBCO-conjugated carboxyrhodamine C110 to achieve specific cell surface labeling (Fig. 1e). Confocal microscopy confirmed successful localization of carboxyrhodamine C110 on the CAR-T cell surface, illustrating efficient and targeted labeling (Fig. 1f).

To optimize labeling concentrations, we conducted a titration series with Ac4ManNAz and carboxyrhodamine C110 ranging from 5 µM to 80 µM (Fig. S3a-c). Flow cytometry analysis revealed that the combination of these compounds achieved efficient fluorescence labeling at all tested concentrations, with 10 µM and 20 µM of each compound yielding fluorescence saturation and yielding maximal labeling efficiency without compromising cell viability. Importantly, increasing concentrations beyond the thresholds did not further enhance signal intensity, indicating a saturation threshold (Fig. S3a-c). Based on these findings, we selected 20 µM Ac4ManNAz and 20 µM carboxyrhodamine C110 as final working concentrations to balance optimal labeling efficiency with minimal cytotoxicity (Fig. S3d). Cytotoxicity assays verified that this labeling protocol did not interfere with CAR-T cell proliferation (Fig. 1g), functional cytokine release, including TNF-α, IFN-γ, Granzyme B (GrzB), and Perforin (Fig. 1h), or transcriptomic profiles as assessed by RNA-seq (Fig. S3e). The fluorescent signal remained detectable for up to 10 days on the proliferating cells (Fig. 1i), demonstrating the durable labeling stability via click-chemistry-mediated glycolabeling. Notably, this click-chemistry labeling approach was successfully applied to various cell types, including NKT cells, NK cells, monocytes, dendritic cells and macrophages (Fig. S3f, g), indicating its broad applicability.

### Enhanced cytotoxicity of CAR-T cells via click chemistry conjugation with MMAE

We utilized click chemistry to conjugate the cytotoxic drug monomethyl auristatin E (MMAE) to the CAR-T cell surface, to achieve two goals: (1) to augment the cytotoxic potential of CA9-targeted CAR-T cells, and (2) to enhance the specificity of MMAE delivery to tumor bed and thereby mitigating systemic toxicity. MMAE has been shown to efficiently kill tumor cells, while CAR-T cells exhibit relatively strong resistance to MMAE after 24 hours of exposure (Fig. S4a, b). Moreover, MMAE treatment did not affect CAR-T functional cytokine release, including IFN-γ, GrzB, and Perforin (Fig. S4c), nor did it largely alter transcriptomic profiles over time (Fig. S4d-f), which supports the feasibility of the CAR-T-D-C approach. Then, we opted for a cleavable valine-citrulline (val-cit) linker^19^, which is susceptible to be cleaved by tumor-secreted Cathepsin B protease (CTSB), known to be overexpressed and activated in a variety of tumors^25^. This linker system, designated as DBCO-PEG4-Val-Cit-PAB-MMAE (DBCO-MMAE), was engineered to facilitate tumor-specific drug release, for enhancing CAR-T-mediated cytotoxicity within the TME (Fig. 2a).

**Figure 2.**
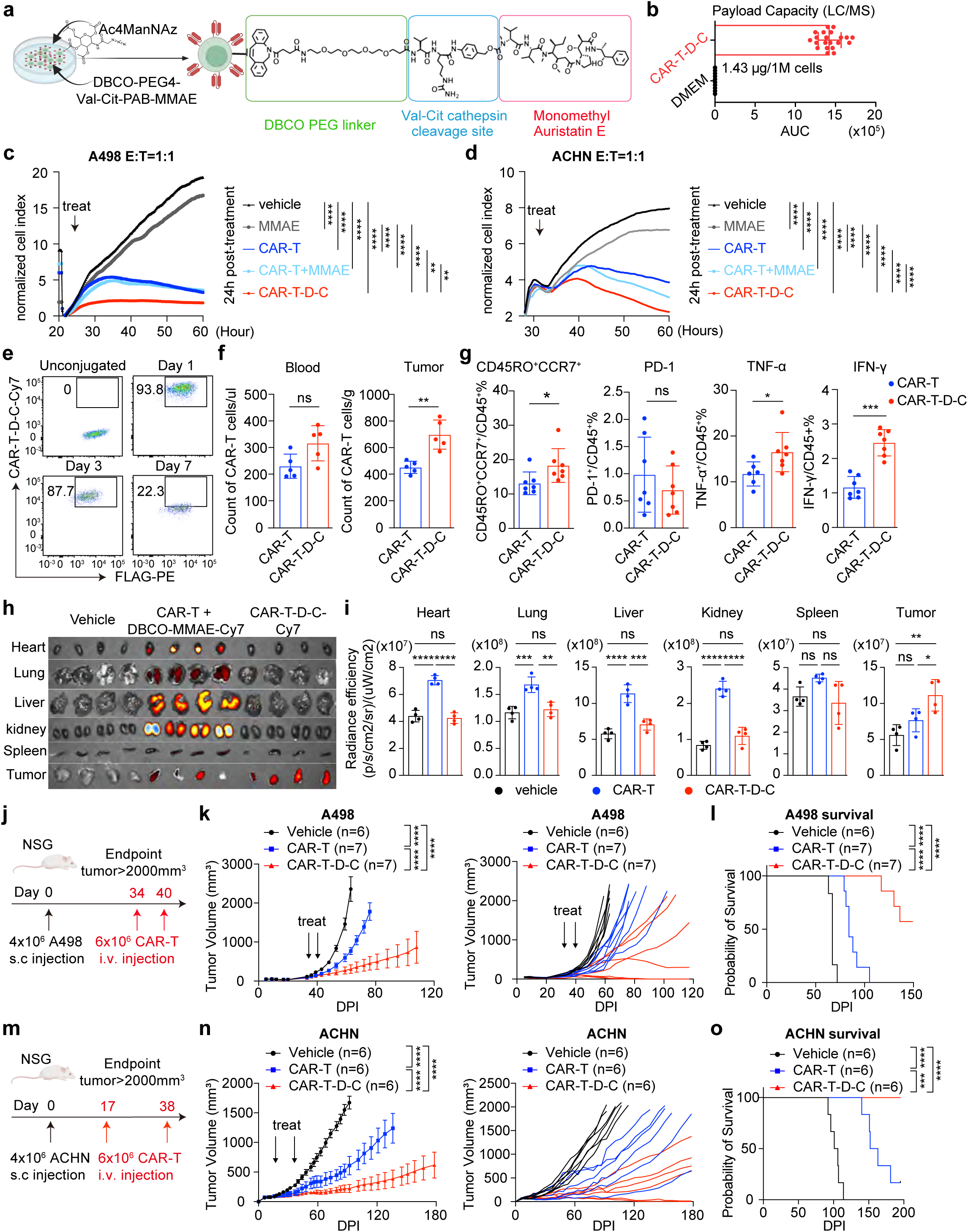
Precise payload delivery and enhanced therapeutic efficacy of anti-CA9 CAR-T-D-C against human kidney cancer cell lines. **(a)** Schematics representation of the structure of DBCO-PEG4-Val-Cit-PAB-MMAE conjugated CAR-T cells. **(b)** Quantification of drug payload carried by CAR-T-D-C using LC/MS. **(c, d)** Real-Time Cell Analysis (RTCA) killing assays were performed to assess CAR-T killing ability against A498 (**c**) or ACHN (**d**). The cancer cells were co-cultured with anti-human CA9 CAR-T cells with or without MMAE conjugation (CAR-T-D-C, CAR-T, respectively), or CAR-T cells with free MMAE (0.25 μM) (CAR-T+MMAE), or free MMAE (0.25 μM), or DMSO vehicle control. E:T: effector to target ratio. Statistical analysis was performed using two-way ANOVA. Statistical significance is denoted as follows: ∗∗p < 0.01; ∗∗∗p < 0.001; ∗∗∗∗p < 0.0001. **(e)** Flow cytometry analysis of surface payload retention on CAR-T-D-C cells in vivo. Peripheral blood was collected from mice via orbital bleed at 1-, 3-, and 7-days post-infusion. **(f)** CAR-T cell counts in blood (left, normalized to volume) and tumors (right, normalized to weight). **(g)** Cell surface marker expression and cytokine release of tumor-infiltrating CAR-T cells isolated on day 10 post-treatment. **(h, i)** IVIS imaging of DBCO-MMAE-Cy7 biodistribution across organs (h) and quantification of MMAE-Cy7 radiance efficiency (i). Tumor-bearing mice were treated with PBS control (vehicle), CAR-T with free DBCO-MMAE-Cy7 (CAR-T + DBCO-MMAE-Cy7), or CAR-T-D-C-Cy7. **(j, k)** In vivo efficacy of anti-CA9 CAR-T cell therapy in A498 kidney cancer xenograft model. Four million A498 cancer cells were subcutaneously injected into NSG mice, followed by 2-times intravenous injection of anti-CA9 CAR-T (n=7), anti-CA9 CAR-T-D-C (n = 7), or PBS vehicle control (n = 6) at the indicated time point. **(l)** Survival curve of mice bearing A498 kidney tumors. **(m, n)** In vivo efficacy of anti-CA9 CAR-T cell therapy in ACHN kidney cancer xenograft model. ACHN cancer cells were subcutaneously injected into NSG mice, followed by 2-times intravenous injection of anti-CA9 CAR-T (n=6), anti-CA9 CAR-T-D-C (n = 6), or PBS vehicle control (n = 6) at the indicated time point. **(o)** Survival curve of mice bearing ACHN kidney tumors. Tumor volume was measured every 3 to 4 days. NSG: NOD.Cg-Prkdc scid Il2rg tm1Wjl /SzJ mice. s.c.: subcutaneous injection. i.v. intravenous injection. Statistical significance was determined using two-way ANOVA. Significance levels are represented as: ∗p < 0.05; ∗∗p < 0.01; ∗∗∗∗p < 0.0001.

To prepare drug conjugated CAR-T cells, we labeled anti-CA9 CAR-T cells with Ac4ManNAz and subsequently incubated with DBCO-MMAE for conjugation. LC-MS analysis confirmed that the val-cit linker was effectively cleaved by cathepsin B present in cancer cell supernatant, enabling MMAE release from the CAR-T-D-C module (Fig. S5a, b). Notably, the addition of exogenous Cathepsin B did not affect cytotoxicity: the MMAE-loaded CAR-T cells (CAR-T-D-C ± CTSB) displayed comparable killing at multiple E:T ratios (1:1, 1:2, 1:5) (Fig. S5d-f). This finding suggests that the Cathepsin B secreted by tumor cells is sufficient to cleave the val-cit linker. We further quantified the drug payload, finding that 1 million CAR-T cells carry approximately 1.43 µg of MMAE (Fig. 2b), a dose well below the reported lethal threshold in mice (1-2 mg/kg).

We evaluated the cytotoxic efficacy of MMAE-conjugated CAR-T cells (defined as MMAE CA9 CAR-T-D-C, or CAR-T-D-C for short) against A498 and ACHN cells using RTCA assay (Fig. 2c, d). For A498 cells, after a 24-hour killing period, the free MMAE (0.25 µM) only slowed cell growth by ∼10%. Conventional CAR-T cells achieved ∼80% killing, and the addition of free MMAE did not further enhance their killing efficacy (CAR-T+MMAE) (Fig. 2c). In contrast, CAR-T-D-C significantly outperformed all other groups in tumor cell killing, achieving ∼92% (Fig. 2c). Flow cytometry analysis after a 3-day coculture revealed that CAR-T-D-C exhibited significantly increased secretion of IFN-γ and TNF-α compared with both the CAR-T and CAR-T+MMAE groups (Fig. S5g).

For ACHN cells, the anti-CA9 CAR-T cell-mediated killing was slower compared to A498 cells, likely due to lower CA9 expression and the inherent harder-to-treat nature of ACHN cells (Fig. 2d). After a 24-hour killing period, free MMAE (0.25 µM) slowed ACHN growth by about 20%. In contrast to A498 cells, CAR-T cells alone achieved ∼50% ACHN cell killing, and the addition of free MMAE further enhanced their killing efficacy by ∼65% (CAR-T+MMAE) (Fig. 2c). Consistent with the results observed with A498 cells, CAR-T-D-C demonstrated superior cytotoxicity, achieving ∼80% killing compared to all other groups (Fig. 2d).

### Localized payload delivery and enhanced anti-tumor efficacy of CAR-T-D-C

To gain a comprehensive understanding of how CAR-T-D-Cs perform both in vitro and in vivo, we initially conjugated the DBCO-MMAE with Cy7 at the MMAE end (Fig. S6a-c). We cocultured A498 with CAR-T-D-C and used time-lapse confocal imaging to track the drug release and trafficking dynamics. Time-lapse confocal imaging revealed that MMAE was gradually released from CAR-T-D-C, preferentially accumulated at the tumor site, and entered tumor cells. This process led to effective tumor cell killing under the combined pressure of CAR-T and the drug payload, while preserving CAR-T-D-C cell viability (Fig. S6d).

Prior to evaluating in vivo efficacy, we assessed the stability of CAR-T-D-Cs in vivo. NSG mice were treated with CAR-T-D-Cs, and blood samples were collected orbitally to monitor CAR-T-D-C retention. CAR-T-D-Cs remained present on the cell surface in vivo through day 7 (Fig. 2e). Subsequently, we treated mice bearing A498 tumors with either CAR-T or CAR-T-D-C cells. Blood and tumor samples were collected to compare the number of CAR-T cells. While there was no significant increase in CAR-T-D-C cells in the blood relative to the CAR-T control, there was a substantial increase in the number of CAR-T-D-C cells within the tumors compared to the CAR-T group (Fig. 2f). Moreover, CAR-T-D-C cells exhibited a higher proportion of CD45RO^+^CCR7^+^ phenotype and increased secretion of TNF-α and IFN-γ, while PD-1 expression remained statistically unchanged (Fig. S7a, Fig. 2g). Notably, the CAR-T-D-C approach delivered MMAE-Cy7 specifically to the tumor site, whereas the nonconjugated drug was systemically distributed across all organs (Fig. 2h-i). Given the potential systemic toxicity of the free drug, all subsequent in vivo experiments compared CAR-T cells with CAR-T-D-C cells exclusively.

To assess therapeutic efficacy in vivo, we utilized xenograft models with cancer cells of human origin. NSG mice were implanted subcutaneously with either A498 (Fig. 2j) or ACHN (Fig. 2m) cells. Treatment groups included a vehicle control, anti-CA9 CAR-T cells, and CAR-T-D-C. Tumor treatment commenced once volumes reached 100 mm³, with 4 million CAR-T cells administered intravenously. In the A498 model, 3 out of 7 mice with CAR-T-D-C treatment achieved complete tumor regression, far surpassing the effectiveness of the two control groups (Fig. 2k). The overall survival of A498 bearing mice was significantly improved, with 3 out of 7 mice completely tumor-free and 1 out of 7 showing controlled tumor growth over a 5-month monitoring period (Fig. 2l). Similarly, in the ACHN model, CAR-T-D-C cells demonstrated superior antitumor efficacy, with 2 out of 6 mice reaching complete tumor remission (Fig. 2n) and overall survival extended by more than 3 months (Fig. 2o). Tumor weights remained largely unaffected during this period (Fig. S7b-c). Our results showed that MMAE CA9 CAR-T-D-C cells are superior to unconjugated anti-CA9 CAR-T cells against kidney cancer cells in culture and in xenograft tumors.

### Application of CAR-T-D-C across multiple tumor types and antigen targets

To demonstrate the versatility of our CAR-T-D-C platform, we aim to expand the strategy to additional cancer types beyond kidney cancer. We selected the HT29 colon cancer cell line for its high surface expression of CA9 (Fig. 3a) and demonstrated that our anti-CA9 scFv clone 1 construct exhibits strong binding affinity to the HT29 cells (Fig. 3b). Consistent with this finding, co-culture of HT29-GFP-luciferase cells with anti-CA9 CAR-T cells resulted in robust killing of the cancer cells across various E:T ratios (1:5, 1:2, 1:1) after 24h incubation (Fig. 3c). Additionally, we assessed the impact of MMAE CA9 CAR-T-D-Cs on HT29 cells through RTCA assay, evaluating the killing efficacy of CAR-T-D-Cs, CAR-T+MMAE, MMAE alone, CAR-T alone and control vehicle group (Fig. 3d). Importantly, the RTCA assay of HT29 cells demonstrated that, similar to kidney cancer cell lines, CAR-T-D-C cells exhibited a significantly enhanced cancer cell-killing effect compared to unconjugated CAR-T cells (Fig. 3d).

**Figure 3.**
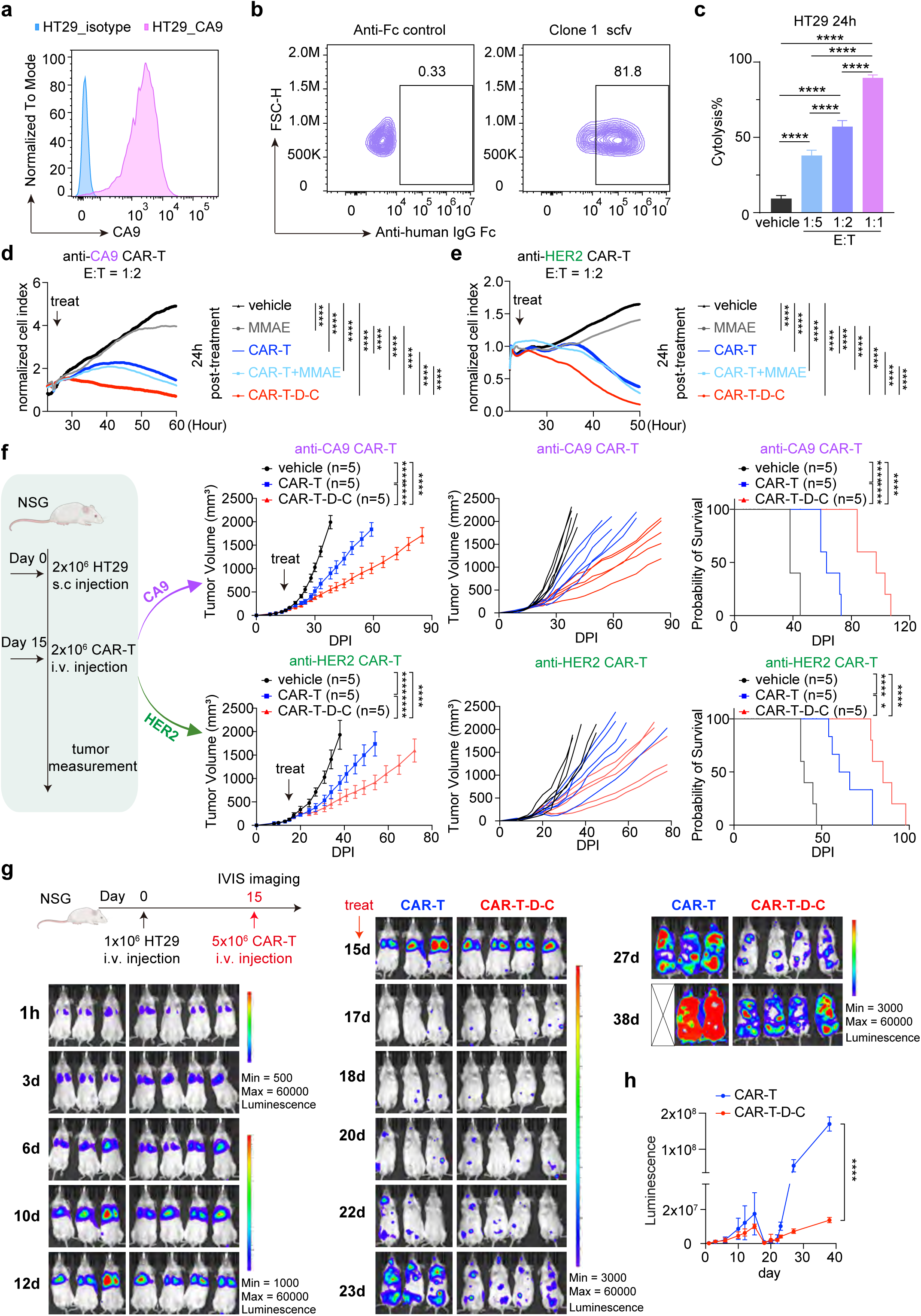
Broader application of CAR-T-D-C beyond kidney cancer to enhance CAR-T cell efficacy. **(a)** Flow cytometry showing the surface CA9 expression level of HT29 cell line. **(b)** Assessment of the binding ability of CA9 scFv clone 1 to HT29 cells using flow cytometry. The scFv was expressed in the vector with human IgG1 heavy chain constant region. **(c)** Coculture assay of HT29 (GFP-Luc) and anti-CA9 CAR-T cells. The cancer cells were co-cultured with anti-human CA9 CAR-T cells at effector to target ratios or with DMSO vehicle control. Cytolysis (%) was calculated by comparing the luminescence of HT29 (GFP-Luc) cells treated with or without CAR-T cell. Cytolysis (%) =100 – Viability (%). **(d, e)** Real-Time Cell Analysis (RTCA) killing assays were applied evaluate the killing efficiency of anti-CA9 CAR-T **(d)** or anti-HER2 CAR-T **(e)** against HT29. HT29 cancer cells were cocultured with different treatments, including free MMAE (0.25 μM), CAR-T alone, CAR-T with 0.25 μM free MMAE (CAR-T+MMAE), CAR-T cells metabolically conjugated with MMAE (CAR-T-D-C), or PBS vehicle control (vehicle). E:T ratio was indicated. Statistical analysis was performed using two-way ANOVA. Statistical significance is denoted as follows: ∗p < 0.05; ∗∗p < 0.01; ∗∗∗p < 0.001; ∗∗∗∗p < 0.0001. **(f)** In vivo efficacy of anti-CA9 CAR-T cells (upper panels) and anti-HER2 CAR-T cells (lower panels) in the HT29 colon cancer xenograft model. Tumor growths were monitored every 3 to 4 days. Five mice were used per group. Statistical significance was assessed using two-way ANOVA. Significance levels: ∗p < 0.05; ∗∗∗p < 0.001; ∗∗∗∗p < 0.0001. **(g, h)** IVIS imaging of tumor progression **(g)** and quantification of in vivo bioluminescence **(h)** in the HT29-GL lung metastasis model.

To further evaluate the applicability of our CAR-T-D-C platform for targeting a range of tumor antigens, we engineered CAR-T cells directed against HER2, a validated antigen overexpressed in HT29 cells (Fig. S8a). Unlike CA9, HER2 undergoes antigen endocytosis upon antibody binding (Fig. S8b). Using lentiviral transduction, we generated anti-HER2 CAR-T cells and evaluated their cytotoxic effects in HT29 cells. Consistent with our findings for anti-CA9 CAR-T cells, HER2 CAR-T-D-C cells showed significantly enhanced tumor cell killing in RTCA assays, outperforming MMAE, CAR-T and CAR-T+MMAE groups (Fig. 3e).

To further confirm therapeutic potential, we conducted in vivo studies in NSG mice bearing subcutaneous HT29 tumors, treated with either anti-CA9 CAR-T or anti-HER2 CAR-T cells (Fig. 3f). In both groups, the CAR-T-D-Cs significantly improved tumor regression compared to both vehicle controls and CAR-T alone. Moreover, the overall survival was prolonged for over one month in the anti-CA9 CAR-T-D-C group and for over three weeks in the anti-HER2 CAR-T-D-C group compared with the corresponding CAR-T groups.

We then assessed the anti-tumor activity using a lung metastasis model in NSG mice. In this model, HT29 cells were injected intravenously (IV) to induce lung metastases. The mice then received a single IV injection of PBS, CAR-T or CAR-T-D-C (Fig. 3g). Mice treated with CAR-T or CAR-T-D-C demonstrated a marked reduction in tumor burden after 2 days of treatment. Although all mice eventually experienced tumor progression, a single infusion of CAR-T-D-C led to a slower progression and prolongation of overall survival (Fig. 3g-h, Fig. S8c).

We performed an additional cytotoxicity assay to compare the killing activity of CAR-T-D-C and the anti-HER2 ADC (trastuzumab vedotin) in the HT29 model. Killing curves of anti-HER2 ADC were generated across a range of concentrations, with viability readouts collected at 24, 48, and 72 hours. The ADC exhibited a dose- and time-dependent cytotoxicity manner. Minimal killing was observed at 24 hours across all concentrations, whereas higher concentrations (0.625–5 µM) induced almost complete cell elimination by 48–72 hours. At 0.3125 µM, the ADC killed approximately 50% of cells at 48 hours and approximately 80% by 72 hours, while lower concentrations produced only modest effects even at later time points (Fig. S8d).

We next performed RTCA killing assays to evaluate CAR-T-D-C in comparison with multiple treatment conditions, including anti-HER2 CAR-T alone, MMAE alone, CAR-T+MMAE, anti-HER2 ADC alone (trastuzumab vedotin), ADC+CAR-T, and the DMSO vehicle control (Fig. S8e). Across all conditions tested, CAR-T-D-C consistently demonstrated superior antitumor activity relative to every control group. The ADC alone, despite delivering an equivalent cytotoxic payload, displayed substantially slower and weaker killing than either CAR-T cells or CAR-T-D-C. Furthermore, the ADC + CAR-T combination showed reduced activity compared with CAR-T alone, likely due to ADC-mediated epitope occupancy that limits CAR-T engagement. These findings indicate that the dual-function CAR-T-D-C strategy outperforms either individual modality or their combinations (Fig. S8e).

### Spatial transcriptomics reveal enhanced CAR-T-D-C function in a lung metastasis model

To gain an unbiased understanding of the status of CAR-T/CAR-T-D-C cells in vivo, we employed a HT29 lung metastasis model and performed spatial transcriptomics to profile the CAR-T localization and functional states within the TME (Fig. 4a). Mice bearing established lung metastases were treated with either CAR-T or CAR-T-D-C, and lungs were harvested 24 h post-treatment for analysis (Fig. 4a, Fig. S9).

**Figure 4.**
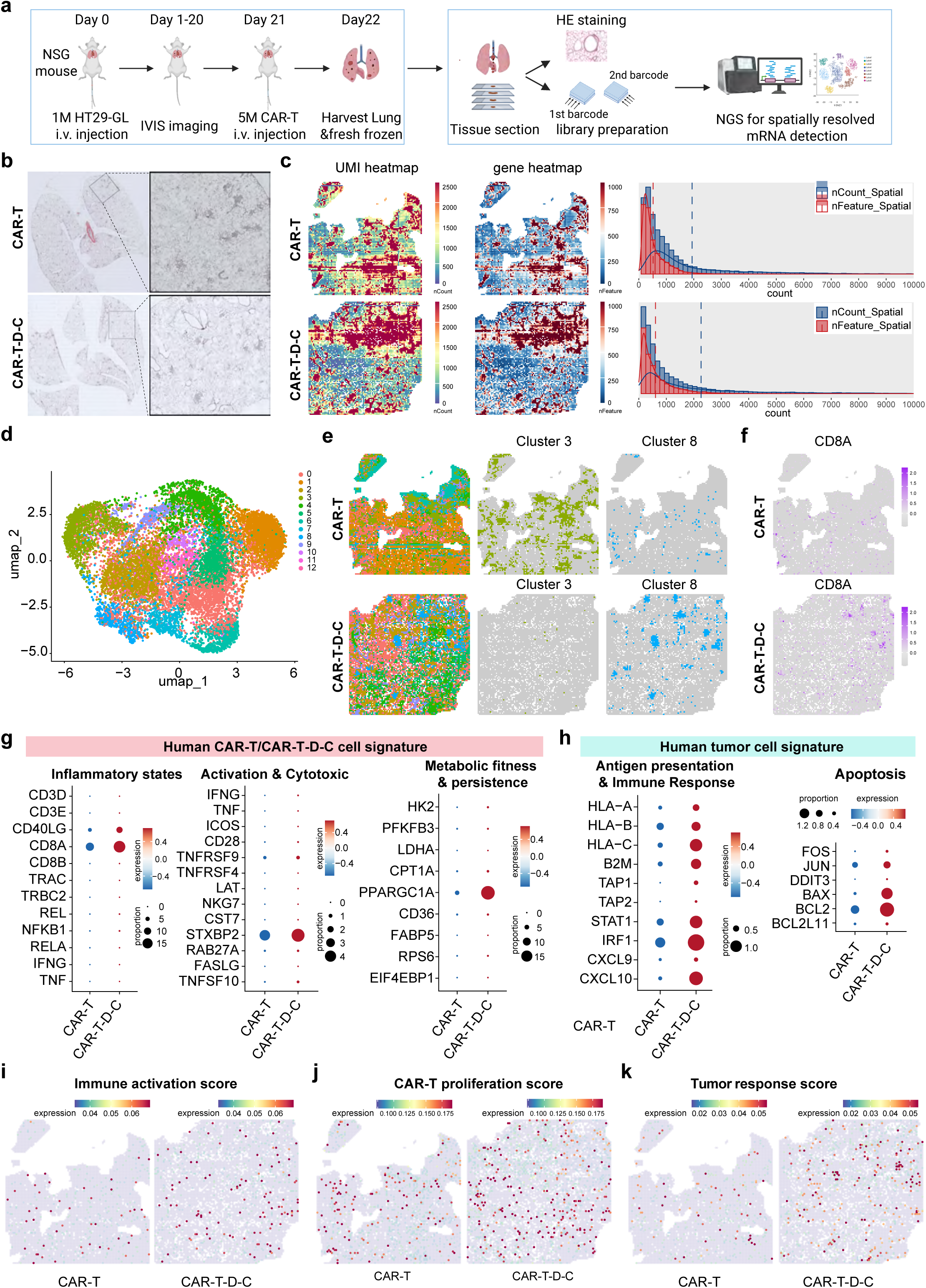
Spatial transcriptomic analysis of CAR-T localization and function in the lung metastasis model. **(a)** Schematic overview of the spatial transcriptomics workflow. **(b)** H&E staining of the lung sections showing metastatic tumor regions. **(c)** Quality control metrics for spatial transcriptomics: total UMI counts per spot and representative UMI heatmap demonstrating uniform coverage across tissue sections. **(d)** UMAP visualization of transcriptionally distinct clusters identified from the gene-by-pixel expression matrix. **(e)** Spatial mapping of clusters 3 (tumor-enriched) and 8 (mixed tumor–immune regions), respectively. **(f)** Feature plots of CD8A expression showing CAR-T cells localization. **(g-h)** Bubble plots depicting representative gene signature of human CAR-T cells **(g)** and human tumor cells **(h)**, with bubble size indicating the proportion of spots expressing each gene and color intensity representing normalized expression levels. **(i-k)** Spatially resolved scores of immune activation score **(i),** CAR-T proliferation score **(j)** and tumor response score **(k)**, calculated from signature gene sets using average expression per spot across the tissue sections.

Hematoxylin and eosin (H&E) staining of lung sections confirmed metastatic tumor regions (Fig. 4b). Data quality was validated by UMI counts per spot and representative UMI heatmaps, which demonstrated uniform coverage across tissue sections (Fig. 4c). A gene expression heatmap further showed that the dataset was partitioned into 12 clusters based on transcriptomic profiles (Fig. S9b). UMAP projection identified these distinct clusters, with cluster 3 corresponding to tumor regions and cluster 8 representing mixed tumor–CAR-T cell regions (Fig. 4d, Fig. 10a-b).

Comparative spatial mapping demonstrated striking differences between the two treatment groups (Fig. 4e-f). CAR-T-D-C cells were predominantly localized within tumor-rich regions, forming mixed CAR-T–tumor areas, whereas CAR-T cells were more diffusely distributed throughout the TME and showed limited colocalization with tumor cells (Fig. 4b, Fig. 4e), indicating that CAR-T-D-C cells had better tumor infiltration. Feature plots of CD8A further revealed a greater abundance and spatial clustering of CAR-T cells in the CAR-T-D-C group compared with the CAR-T group. These cells were preferentially localized within tumor regions, highlighting the enhanced tumor infiltration achieved by CAR-T-D-C therapy (Fig. 4f).

Functionally, CAR-T-D-C treatment showed markedly elevated expression of key effector molecules. These included markers of inflammatory states, immune activation and cytotoxic phenotype, as well as indicators of metabolic fitness and persistence (Fig. 4g), consistent with the enhanced therapeutic efficacy. Moreover, CAR-T-D-C treated tumors exhibited stronger antigen presentation signatures, increased immune response and more pronounced apoptotic signatures (Fig. 4h). The CAR-T-D-C treatment was also associated with improved immune activation scores, higher CAR-T proliferation score, as well as increased tumor response score compared with CAR-T treatment (Fig. 4i-k).

### CAR-T-D-C reshapes tumor microenvironment by enhancing antigen presenting cells

To further study how the CAR-T-D-C affects the TME and anti-tumor immunity, we utilized the Colon26 syngeneic tumor model with adoptive transfer of CAR-Ts of mouse origin, which unlike xenograft models, provides a fully immunocompetent setting for the study of tumor immunology (Fig. 5a). First, we engineered the Colon26 cell line to overexpress human CA9 using lentiviral transduction (Fig. 5b) and implanted these cells subcutaneously into BALB/c mice following the experimental timeline outlined in Fig. 5c. Once the tumors reached an approximate volume of 100 mm³, we initiated treatment by administering vehicle control (PBS), anti-CA9 mouse CAR-T cells, or anti-CA9 mouse CAR-T-D-Cs. Our in vivo experiments demonstrated that the CAR-T-D-Cs exhibited significantly enhanced therapeutic efficacy compared to the vehicle control and CAR-T groups. In the CAR-T group, 2 out of 7 mice achieved complete tumor regression, whereas in the CAR-T-D-C group 6 out of 8 mice achieved complete tumor regression (Fig. 5c). Moreover, the CAR-T-D-C group showed significant improved overall survival among all three groups (Fig. 5d).

**Figure 5.**
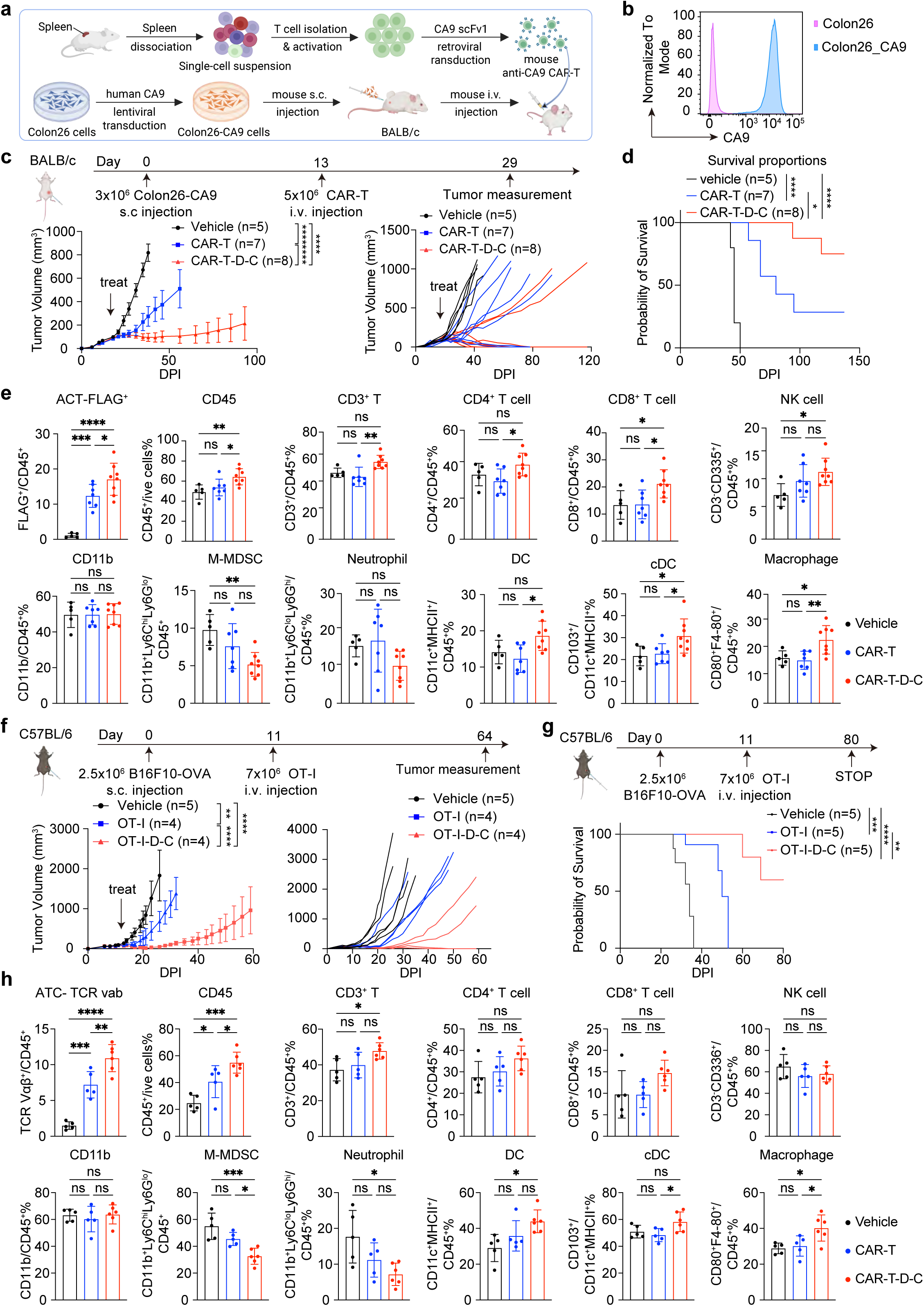
CAR-T-D-C reshapes the tumor microenvironment within solid tumors. **(a)** Schematic illustration of the mouse anti-CA9 CAR-T generation and treatment strategy in the syngeneic Colon26-CA9 colon cancer model in BALB/c mice. **(b)** Flow cytometry showing the surface CA9 expression level of Colon26 cell line with or without human CA9 overexpression. **(c)** Upper panel: Schematic representation of the CAR-T treatment strategy in the syngeneic Colon26 colon cancer model in BALB/c mice. Lower panel: Growth curves showing tumor volume measured over time under different treatments: conventional CA9 CAR-T (n = 7), MMAE CA9 CAR-T-D-C (n = 8), or PBS vehicle control (n = 5). Data are presented as mean ± SEM. Statistical comparisons were performed using two-way ANOVA for each group against PBS vehicle control group. Statistical significance is denoted as follows: ∗p < 0.05; ∗∗∗p < 0.001; ∗∗∗∗p < 0.0001. **(d)** Overall survival analysis of Colon26-CA9 tumors in BALB/c mice following treatments with PBS, CAR-T and CAR-T-D-C. Statistical comparisons were made using the Log-rank (Mantel-Cox) test for each treatment group against the vehicle control group. Statistical significance is denoted as follows: ∗p < 0.05; ∗∗∗p < 0.001; ∗∗∗∗p < 0.0001. **(e)** Tumor-infiltrating immune cell composition in Colon26-CA9 tumors following treatments with PBS vehicle control, CAR-T and CAR-T-D-C. Statistical comparisons were made with one-way ANOVA. Statistical significance is denoted as follows: ∗p < 0.05; ∗∗∗p < 0.001; ∗∗∗∗p < 0.0001. **(f)** Upper panel: Schematic representation of the OT-I T cell treatment strategy in the syngeneic mouse B16F10-OVA melanoma model. Lower panel: Growth curves of B16F10-OVA tumors in C57BL/6 mice following different treatments: OT-I T cells (n = 4), OT-I T-D-C (n = 4), or PBS vehicle control (n = 5). Data are presented as mean ± SEM. Statistical comparisons were performed using two-way ANOVA for each group against PBS. Statistical significance is denoted as follows: ∗p < 0.05; ∗∗∗p < 0.001; ∗∗∗∗p < 0.0001. **(g)** Overall survival analysis of the B16F10-OVA tumor model treated with OT-I T cells, OT-1-D-Cs, or PBS vehicle control. Statistical comparisons were made using the Log-rank (Mantel-Cox) test for each treatment group against the vehicle control group. Statistical significance is denoted as follows: ∗∗p < 0.01; ∗∗∗p < 0.001. **(h)** Tumor-infiltrating immune cell composition in B16F10 tumors following treatments with PBS vehicle control, CAR-T and CAR-T-D-C. Statistical comparisons were made with one-way ANOVA. Statistical significance is denoted as follows: ∗p < 0.05; ∗∗∗p < 0.001; ∗∗∗∗p < 0.0001.

To dissect the immune landscape within the TME following treatment, we performed comprehensive flow cytometry analysis of tumor-infiltrating immune cells (Fig. S11). We observed a significant increase of adoptively transferred FLAG^+^ cells in the CAR-T-D-C group compared with the CAR-T group (Fig. 5e). Additionally, the data revealed a substantial increase in the populations of CD45^+^ immune cells, CD8^+^ T cells, CD11b^+^F4/80^+^ macrophages and CD103^+^CD11c^+^MHCII^+^ conventional dendritic cells within the tumors of mice treated with the CAR-T-D-C compared to both vehicle and CAR-T only groups (Fig. 5e). This suggests that the enhanced therapeutic effect may be attributed, in part, to improved antigen presentation and subsequent activation of the immune response. Although CD3^+^ T cells and CD4^+^ T cells were significantly increased in the CAR-T-D-C group compared with the CAR-T group, they were not significantly different from the PBS control group. However, there was still an observable increasing trend. The CD11b^+^ monocytes and the CD11b^+^Ly6C^lo^Ly6G^hi^ Neutrophil population remained relatively unchanged among groups, while the immunosuppressive CD11b^+^Ly6C^hi^Ly6G^lo^ M-MDSC population is markedly decreased in the CAR-T-D-C group compared with the vehicle control group (Fig. 5e).

To test the breath of applicability of CAR-T-D-C, we extended the application of the click-chemistry-conjugated MMAE beyond CAR-T cells to other engineered T cell modalities, such as TCR-transgenic T cells. Specifically, we applied this DBCO-MMAE conjugate with the mouse OT-I T cells, which express TCR that recognize ovalbumin (OVA) residues 257-264 (SIINFEKL) in the context of H2Kb. This approach was tested in a B16F10-OVA melanoma mouse model, a “cold tumor” known for its resistance to conventional checkpoint blockade immunotherapies^26^. Treatment efficacy was compared among vehicle, OT-I T cells alone, and OT-I cells conjugated with MMAE (MMAE OT-I-D-C) (Fig. 5f). The OT-I-D-C group significantly reduced tumor burden, effectively halting tumor progression for over a month and markedly extending survival relative to control groups with 2 out of 4 mice achieved complete tumor regression (Fig. 5g).

To understand the impact of OT-I-D-C therapy on the TME, we conducted comprehensive flow cytometry analysis to characterize tumor-infiltrating immune cells in the B16F10 tumors post-treatment. There was a notable increase in adoptively transferred TCR Vαβ^+^ OT-I cells in the OT-I-D-C group compared to the OT-I group (Fig. 5h). Treatment with OT-I-D-C led to a robust infiltration of CD45^+^ immune cells and significant enrichment in antigen presenting cells (APCs), such as CD11b^+^F4/80^+^ macrophage and CD11c^+^MHCII^+^ dendritic cell (DC) populations compared to both vehicle and OT-I only groups. Additionally, there was an observable trend of increasing CD3^+^ T cells, CD4^+^ T cells, and CD8^+^ T cells in the OT-I-D-C group (Fig. 5h).

These findings indicate that both CAR-T-D-C and OT-I-D-C not only directly target tumor cells but also reshapes the tumor microenvironment by facilitating the infiltration of immune cells, including T cells and key APCs, promoting an adaptive immune response that may enhance overall therapeutic efficacy (Fig. 5e, h).

### CAR-T-D-C triggers a systemic antitumor immune response

Chemotherapy-induced antigen spreading is well-documented in enhancing immune surveillance through the antigen spreading^8, 27^. Based on this mechanism, we hypothesized that the CAR-T-D-C could similarly stimulate systemic antitumor immunity, as APC-mediated acquisition and display of tumor antigens is crucial for initiating a robust T-cell response across distant tumor sites. To test this hypothesis, we assessed whether the CAR-T-D-C therapy could elicit a systemic antitumor immune response in syngeneic models bearing bilateral tumors. We implanted Colon26 tumors bilaterally in the flanks of mice, with one tumor overexpressing the target antigen (1° Colon26-CA9) and the other lacking it (2° Colon26). Following the experimental timelines outlined in Fig. 6a, we administered mouse CA9 CAR-T cells or mouse MMAE CA9 CAR-T-D-C intravenously. Notably, in the Colon26 model, treatment with both CAR-T-D-C and CAR-T cells significantly slowed progression of both the antigen-positive primary (1°) and antigen-negative secondary tumors (2°), with CAR-T-D-C outperformed the unmodified CAR-T group (Fig. 6b-d). 3/8 of the mice reached complete tumor regression even on the antigen-negative side, with a significantly improved overall survival (Fig. 6e). In the bilateral B16F10-OVA and B16F10 model, the OT-I-D-C strongly suppressed tumor progression, eradicating most of the OVA-positive primary tumors and slowing down the growth of OVA-negative secondary tumors, again implying a systemic immune response (Fig. 6h-j). The OT-I-D-C group effectively halt tumor progression for over six weeks and markedly extended overall survival relative to control groups (Fig. 6k).

**Figure 6.**
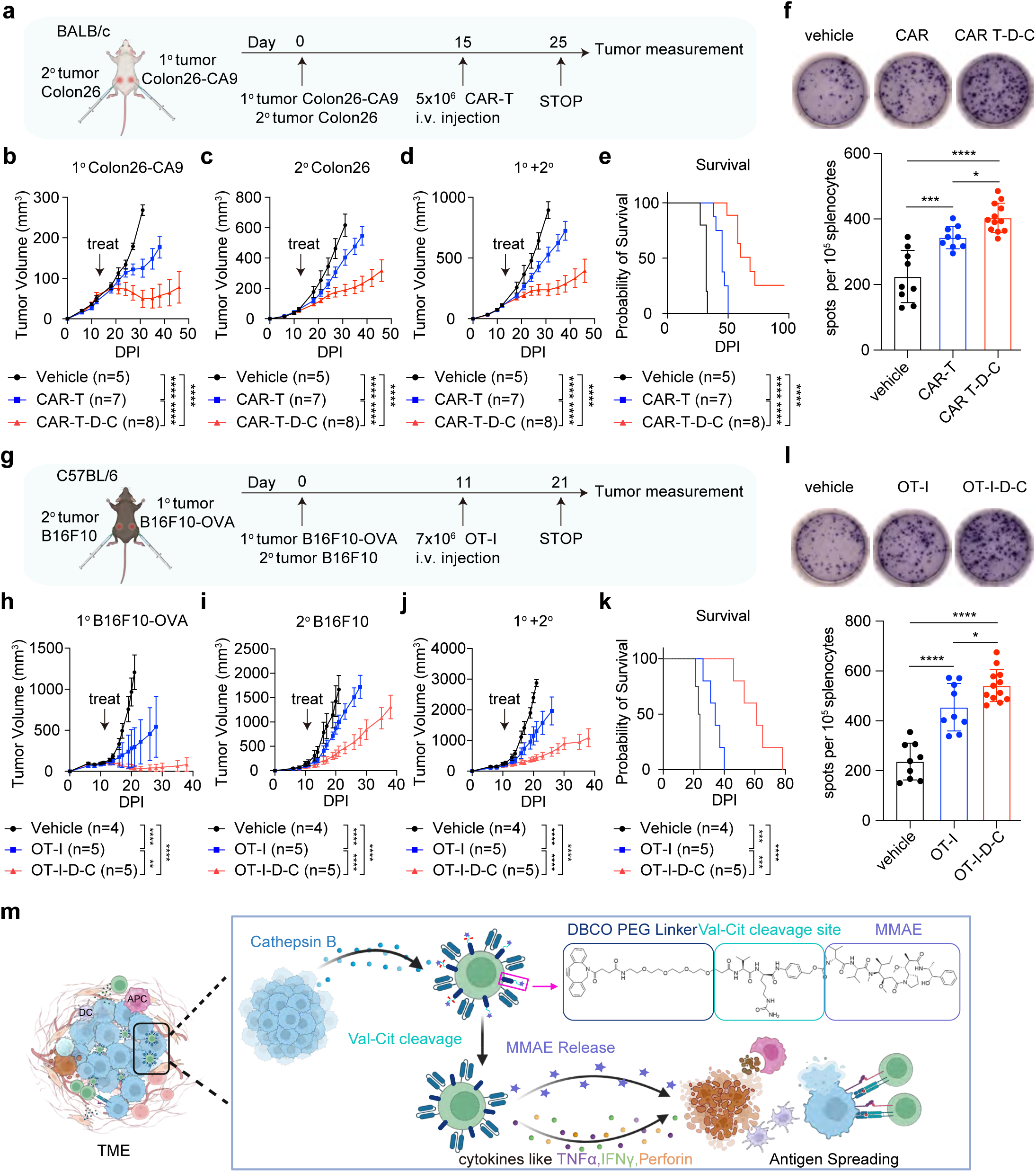
CAR-T-D-C treatment enhances systemic anti-tumor immune response. **(a)** Schematics representation the CAR-T-D-C treatment strategy in a dual tumor model of mouse Colon26 colon cancer to study the systemic anti-tumor response. **(b-d)** Growth curve showing Colon26 tumor volume measured overtime following treatments with PBS vehicle control, CAR-T and CAR-T-D-C. (**b**) Primary tumor Colon26-CA9 with human CA9 overexpression; (**c**) Secondary tumor Colon26 without human CA9 antigen. **(d)** Combined analysis of primary and secondary tumors. Statistical comparisons were made using two-way ANOVA. Statistical significance is denoted as follows: ∗p < 0.05; ∗∗∗∗p < 0.0001. **(e)** Survival of mice with bilateral Colon26-CA9 (primary) and Colon26 (secondary) tumors following CAR-T, CAR-T-D-C, or PBS vehicle control treatment. **(f)** IFN-γ secretion ability of the splenocytes isolated from tumor bearing-mice treated with CAR-T, CAR-T-D-C or PBS vehicle, measured by ELISPOT assay. Statistical comparisons were made with one-way ANOVA. Statistical significance is denoted as follows: ∗p < 0.05; ∗∗∗p < 0.001; ∗∗∗∗p < 0.0001. **(g)** Schematics representation of the OT-I T cell treatment strategy in a dual tumor model with B16F10-OVA (primary) and B16F10 (secondary) in C57BL/6 mice. **(h-j)** Growth curves of the tumor volume measured overtime with indicated treatments. (**f**) Primary tumor with the OVA antigen; (**g**) Secondary tumor without OVA antigen. Statistical comparisons were made using two-way ANOVA. (**j**) Combined analysis of primary and secondary tumors. Statistical significance is denoted as follows: ∗p < 0.05; ∗∗∗∗p < 0.0001. (**k**) Survival of mice with bilateral B16F10- OVA (primary) and B16F10 (secondary) tumors following OT-I, OT-I-D-C, or PBS vehicle treatment. **(l)** IFN-γ secretion ability of the splenocytes isolated from B16F10 tumor bearing-mice treated with OT1-T, OT-I-D-C or PBS vehicle, measured by IFN-γ ELISPOT assay. Statistical comparisons were made using one-way ANOVA. Statistical significance is denoted as follows: ∗p < 0.05; ∗∗∗∗p < 0.0001. (**m**) Schematic summarizing the mechanism of the dual-function CAR-T-D-C approach.

To further study whether CAR-T-D-C induced systemic immune response, we evaluated the activation of tumor-specific endogenous T cells capable of producing IFN-γ via ELISPOT. We isolated splenocytes from PBS, CAR-T and CAR-T-D-C treated mice bearing Colon26-CA9 and restimulated the splenocytes ex vivo with antigen-positive tumor cells Colon26-CA9. As shown in Fig. 6f, re-stimulated splenocytes from CAR-T-D-C treated mice revealed significantly enhanced IFN-γ secretion compared to the controls, confirming that CAR-T-D-C treatment amplified tumor-specific T-cell responses. Similarly, re-stimulation of splenocytes from B16F10-OVA bearing mice demonstrated that OT-I-D-C treatment yielded higher IFN-γ levels compared with vehicle and OT-I only group. These results suggest that CAR-T-D-C and OT-I-D-C therapies induced systemic anti-tumor immunity as reflected in the activation and expansion of tumor-specific T cells capable of targeting tumor sites beyond the primary tumor (Fig. 6l). The dual-function CAR-T-D-C approach precisely delivers therapeutic payloads into the tumor bed, significantly enhances immune cell infiltration, promotes antigen spreading, amplifies systemic immune responses, and ultimately improves anti-tumor immunity (Fig. 6m).

## Discussion

In this study, we developed and validated the CAR-T-D-C as a novel, dual function approach for targeting solid tumors. This approach leverages a cleavable val-cit linker responsive to Cathepsin B to ensure the controlled release and precise accumulation of drug payload within the TME, thereby boosting CAR-T cell efficacy while minimizing systemic toxicity. Our data demonstrated robust anti-tumor efficacy of CAR-T-D-C across various solid tumor models, showcasing both direct antitumor effects and the capacity to stimulate systemic immune responses. These findings offer a different strategy for overcoming the challenges of solid tumor immunotherapy.

Several previous studies have demonstrated different strategies to enhance cell-based cancer therapies using nanoparticle or polymeric micropatches. For examples: Huang et al. employed nanoparticle-carrying T cells to deliver chemotherapy to disseminated tumors rather than solid tumors, and their platform likewise lacked a defined tumor-specific delivery design^28^. Kumbhojkar et al. polarized neutrophils toward an antitumor phenotype with polymeric micropatches to activate NK and T cells for immune response^29^. Yang et al. exploited macrophages as drug carriers by attaching doxorubicin-loaded liposomes via biotin–avidin interactions. This macrophage–liposome system enhanced drug accumulation and penetration in tumors^30^. These studies primarily use immune cells as carriers to transport therapeutic payloads for passive killing effect, but with limited utility of engineered CARs. Tang et al. engineered protein nanogels (NGs) that ‘backpack’ cytokines onto T cells and release them in response to TCR signaling a strategy designed to localize cytokine support for T-cell function^31^. In contrast to the conjugation strategies used in these systems, our platform employs click chemistry to stably and bioorthogonally link CAR-T cells with a cleavable cytotoxic payload and uniquely integrates dual functions: 1) immunotherapy mediated by antigen-specific CAR-T cytotoxicity, and 2) chemotherapy initiated by tumor-localized therapeutic payload release, which resulting in a synergistic antitumor effect.

Click chemistry has shown significant utility in cancer research, particularly in cell engineering^32, 33^. For instance, Lamoot et al. represents an advance in applying click chemistry to attach lipid nanoparticles (LNPs) and Jurkat cells^34^, however, Jurkat cells are not primary T cells nor CAR-Ts. Li et al. further applied click chemistry for efficient, controllable conjugation of lipid nanoparticles (LNPs) to T cells, enabling drug loading^35^, however, this approach does not allow TME-specific cytotoxic payload to be release for potent and specific cancer killing in the tumor. These usage of click chemistry served primarily as a bioorthogonal surface-labeling tool for immune cells with LNPs, without demonstration of therapeutic function. In contrast, our approach transforms click chemistry into a therapeutically functional interface for cancer therapy, enabling coordinated CAR-T–mediated killing together with tumor-restricted drug release.

While CAR-T therapy has revolutionized treatment of hematologic malignancies, its efficacy in solid tumors remains limited by poor tumor infiltration, an immunosuppressive microenvironment, and antigen heterogeneity or loss. Although ADCs are potent targeted therapies, their effectiveness in solid tumors is restricted by the need for efficient antigen internalization, antigen heterogeneity, and limited tissue penetration. ADC is dependent on the endocytosis process for the small molecule payload to internalized into a target cell to kill cancer cells, however, it will face challenges in targets where endocytosis of the ADC-target complex in not efficient. On the contrary, CAR-T-D-C does not require the biologics (CAR) to be internalized via endocytosis, because (1) CAR can recognize the antigen and the killing of cancer cell is in part mediated by the T cells; (2) the chemotherapeutic payload gets released in the presence of enzymes like cathepsin B in the TME, for which the free drug itself is only locally concentrated in the vicinity of tumor and kills cancer cells.

The CAR-T-D-C approach that integrates immunotherapy and chemotherapy is well-suited to address challenges faced by conventional CAR-T therapy and ADC targeted therapy. It transforms CAR-T cell from a conventional effector into a programmable therapeutic node that senses the tumor’s cathepsin-rich microenvironment and responds by locally releasing a cytotoxic payload. Notably, the dual-function CAR-T-D-C strategy achieves superior activity compared with either individual modality (CAR-T alone or ADC alone) or their combinations.

Earlier work by Pan et al. uses azido-sugar labeling of CAR-T cells and artificial BCN labeling of tumor cells to create a synthetic ligand–receptor interaction^36^; however, this study relies on artificial introduction of BCN onto tumor cells, which does not reflect a clinically-viable treatment setting. Zhao et al. employs click chemistry to attach an anti–PD-1 antibody to CAR-T cells, harnessing the low-pH tumor milieu to augment checkpoint blockade activity, but it does not introduce an immuno-chemo dual-function cytotoxic mechanism^37^. Depending on the solid tumor context, CAR-T-D-C can initiate with CAR-T mediated cytotoxicity followed by payload release that amplifies systemic antitumor responses, or with early payload release into TME that promotes antigen spreading and enhances immune activation. CAR-T-D-C, the combinatorial therapeutic approach, establishes a positive feedback loop that elicits a potent systemic immune response plus TME-specific cytotoxic payload release.

The simplicity and versatility of the click labeling process make the CAR-T-D-C approach adaptable to a range of tumor-associated antigens, including human anti-CA9 CAR-T, human anti-HER2 CAR-T and mouse OT-I transgenic TCR-T targeting a model antigen OVA. Integration of drug conjugation into CAR-T cell production via metabolic processes^38^ enhances the efficiency and consistency of adoptive cell transfer (ACT) therapies without increasing manufacturing complexity. the simplicity and versatility of the click labeling process make it an adaptable tool, allowing the CAR-T-D-C approach to be potentially applied across a variety of cell therapies.

A notable advantage of the CAR-T-D-C strategy lies in its ability to transform the immunosuppressive TME into an ally. Solid tumors present a complex array of challenges for CAR-T cell therapy, including limited tumor penetration, immunosuppressive TMEs, and antigenic variability^8^. Instead of merely overcoming the immunosuppressive TME, CAR-T-D-C arms with DBCO-val-cit-MMAE and leverages TME-secreted cathepsin B to achieve precise antigen recognition and localized drug delivery. This approach results in cumulative cytolytic activity mediated by combining the direct tumor-killing effects of CAR-T cells with the locally released drug payload, resulting in enhanced infiltration of adoptively transferred cells, increased tumor antigen release, and robust systemic immune activation. This secondary immune activation is particularly important for overcoming tumor heterogeneity and antigen-loss–driven resistance, a major limitation of conventional CAR-T therapy^39,40, 41^. With these advantages, the CAR-T-D-C approach provides precise control over payload dosing, eliminating the need for repeated administrations and thereby minimizing the risks of drug accumulation and systemic toxicity.

Another key advantage of CAR-T-D-C is its flexibility, where both drug payload and linker chemistry can be customized to exploit the specific metabolic and enzymatic characteristics of the target tumor^19, 42^. This modular design enables the CAR-T-D-C strategy to be tailored to a wide spectrum of tumors with distinct metabolic or enzymatic profiles. Furthermore, the simplicity of the bioorthogonal click chemistry reaction allows the drug conjugation step to be seamlessly incorporated into CAR-T cell manufacturing, without substantially increasing the complexity, duration, or regulatory burden of adoptive cell transfer (ACT) therapies. This integration supports scalable production while maintaining the functionality and viability of the engineered T cells, facilitating translation into clinical applications.

Our study has limitations that can be addressed in future research. First, potential off-target effects associated with CA9 antigen targeting could impact normal tissues, necessitating meticulous evaluation and safety measures. In future experiments, scFvs can be screened and optimized to minimize the off-target effect. Second, the exploration of alternative linker chemistries and drug types tailored to different tumor profiles could further refine and optimize the CAR-T-D-C platform. Third, beyond CAR-T therapy, the applicability of the CAR-T-D-C therapeutic efficacy could be broadened by testing different cell therapy types. Lastly, human PDXs can be tested to further validate the CAR-T-D-C approach in additional clinically relevant models.

In summary, the CAR-T-D-C platform, with its adaptability to multiple cell types, various antigens, and different payloads, represents a 2-in-1 immunochemotherapy for the treatment of diverse cancer types, particularly solid tumors.

## Author Contributions

Conceptualization: SC, SX. Design: SX, YF, FZ, SC. Experiment lead: SX, YF, FZ. Assistance and support: JY, CD, BT, KT, QW, BC, RP, NV, CL, KS, XL, DP, HK, JMC, RF, JZ. Manuscript prep: SX, YF, FZ, SC. Supervision and funding: SC.

## Acknowledgments

We thank Nikolaus Joerg at the Yale west campus imaging core for assistance with imaging. We thank all members in Chen laboratory, as well as various colleagues in Yale Genetics, SBI, CSBC, MCGD, Immunobiology, BBS, YCC, YSCC, and CBDS for assistance and/or discussions. We thank various Yale Core Facilities such as YARC, WCFC, YCGA, HPC, and WCAC for technical support.

SC is supported by fundings from NIH/NCI (R33CA281702), DoD (W81XWH-20-1-0072, W81XWH-21-1-0514, HT94252310472, HT94252510589), Cancer Research Institute (Cancer Research Institute Lloyd J. Old STAR Award (CRI4964) and CLIP KCA Award (CRI13945), Pershing Square Sohn Cancer Research Alliance and Yale Skin SPORE pilot. NV is supported by American Board of Radiology’s B. Leonard Holman Research Pathway Fellowship and the ASTRO Radiation Oncology Seed Grant.

## Methods

### Institutional approval

This study has received institutional regulatory approval. All recombinant DNA and biosafety work were conducted in accordance with the guidelines established by the Yale Environment, Health and Safety (EHS) Committee, under the approved protocol (Chen-rDNA-15-45). Additionally, all animal work was approved by Yale University’s Institutional Animal Care and Use Committee (IACUC) and performed in compliance with approved protocols (Chen protocol# 20068).

### Cell culture

Expi293F, HEK293FT, Lenti-X 293T, A498, ACHN, HT29, Colon26, and B16F10 cell lines were obtained from commercial sources (ThermoFisher, American Type Culture Collection (ATCC), Takara) and were periodically tested negative for mycoplasma contamination. HEK293FT, Lenti-X 293T, HT29 and B16F10 cells were cultured in DMEM (Gibco) media supplemented with 10% FBS (Sigma) and 200 U/mL penicillin–streptomycin (Gibco) (hereafter referred to as D10 medium). A498 and ACHN were cultured in EMEM, with 10% FBS, 200 U/mL penicillin–streptomycin. Colon26 cells were cultured in RPMI-1640 (Gibco) media supplemented with 10% FBS and 200 U/mL penicillin–streptomycin (hereafter referred to as R10 medium). Expi293F cells were cultured in Expi293™ Expression Medium (Gibco). All cell lines used in this paper were maintained at 37°C with 5% CO2.

### Mice

All animal procedures were performed in compliance with protocols approved by the Yale IACUC. Mice were housed with unrestricted access to food and water in a temperature-controlled (approximately 22°C) and humidity-regulated colony room, maintained on a 12-hour light/dark cycle (lights on from 07:00 to 19:00). Daily health assessments were conducted to monitor well-being. Mice aged 8-12 weeks were used for experiments. ATK mice, NOD-scid IL2Rgammanull (NSG) mice, BALB/c mice, C57BL/6 mice and OT-I TCR transgenic mice were used in this study.

### Generation of CA9 CAR binder

The development of CA9 CAR binder was performed as described before^22, 23^.CA9-mRNA were synthesized with the Hiscribe™ T7 ARCA mRNA Kit (with tailing) (NEB, Cat. E2060S) and formulated using the NanoAssemblr® Ignite™ instrument (Precision Nanosystems) according to manufacturers’ guidelines. The particle size of LNP-mRNA was determined using dynamic light scattering (DLS) device (DynaPro NanoStar, Wyatt, WDPN-06). The encapsulation efficiency and mRNA concentration were quantified by Quant-iT™ RiboGreen™ RNA Assay (Thermo Fisher, Cat. R11490). Humanized ATX-GK mice (Alloy Therapeutics), engineered with with human IgG and IgK transgene knock-ins, were immunized with 10μg of CA9 LNP-mRNA per dose for both prime and booster injections on day 0 and day 14, respectively. Blood samples were collected retro-orbitally on days 14 and 21 post-immunization to evaluate the immune response using ELISA. Spleen and lymph nodes from immunized mice were harvested on day 23 for memory B cells isolation using a magnetic positive selection kit (Miltenyi Biotec, Cat. 130-095-838). Approximately 10,000 memory B cells were loaded on a Chromium Next GEM Chip K for single-cell lysis and cDNA first-strand synthesis according to the manufacturer’s protocol, flowed by cDNA amplification. To construct BCR repertoire libraries, 2 μL of amplified cDNA was subjected to two rounds of target enrichment using nested custom primer pairs specific for BCR constant regions. The library was profiled and quantified using the D1000 ScreenTape assay (Agilent) for the TapeStation system and sequenced by paired-end sequencing (26 x 91 bp) on an Illumina MiSeq, aiming to identify CA9-reactive BCRs with a targeted sequencing depth of 5,000 read pairs per cell. Single-cell BCR-seq data were converted to demultiplexed FASTQ files by the Illumina Miseq controller. Data processing was conducted by using Cell Ranger v6.0.1, with reads aligned to a customized V(D)J reference comprising mouse constant genes and human V(D)J genes. The top-enriched clonotype consensus sequences were identified using Loupe V(D)J Browser, and selected sequences were synthesized for subsequent molecular cloning and recombinant antibody expression.

### Molecular cloning

Plasmids were constructed using standard Gibson assembly method. To reconstruct full IgG1 antibodies, the paired heavy- and kappa-variable fragments from top-ranked enriched IgG clones were synthesized as gBlocks and subcloned into pFuse-hIgG1 and pFuse-hKappa vectors (InvivoGen), respectively, for mammalian expression as described previously ^23^. For scFv constructs generation, paired heavy- and kappa-variable fragments were chained via a flexible linker peptide, (GGGGS)_3_, by PCR amplification and subcloned into a pFuse-hIgG1 vector for mammalian expression.

### Antibody production and binding validation by cell surface staining

Monoclonal antibodies (mAbs) were produced by transiently transfection of plasmids into Expi293F cells with either equal amounts of paired heavy- and light-chain expression vectors, or an scFv expression vector, using the ExpiFectamine 293 transfection kit according to the manufacturer’s protocol (ThermoFisher). Five days post-transfection, supernatants containing the secreted mAbs were collected and used as primary antibodies in cell staining assay to determine antibody binding on cell surface of CA9-overexpression cells. For flow cytometry staining, 200 μL of each transfected culture supernatant was incubated with CA9-over-expressed cells for 1 h at 4°C. After incubation, the cells were washed twice and then incubated with anti-human IgG-PE secondary antibodies for an additional 30 min on ice. Binding activity was determined using flow cytometry.

### Human T cell culture isolation

Human CD3 T cells were purified from human PBMCs using Human pan T cell kit (Miltenyi, Cat. 130-096-535). Fresh isolated human CD3 T cells were stimulated with anti-CD3/CD28 dynabeads (ThermoFisher, Cat. 11161D) for 48 h and cultured at a concentration of 1-2 × 10^6^ cells/ml in X-VIVO 15 media (Lonza) supplemented with 5% Human AB serum + IL-2 (100 U/ml).

### Mouse T cell isolation and culture

Spleens were isolated from indicated mouse strains and placed in ice-cold washing buffer (PBS with 2% FBS). Spleens were then dissociated by mashing through a 100 μm filter in washing buffer. Red blood cells were lysed with ACK lysis buffer (Lonza), incubated for 2 mins at room temperature, and washed with washing buffer. The resulting cell suspensions were filtered through a 40 μm filter and resuspended with MACS Buffer (PBS + 0.5% BSA + 2 μM EDTA). Pan T cells from BALB/c mice or CD8 T cells from OT-I mice were isolated using the Miltenyi Pan T Cell Isolation Kit II (mouse, Cat. 130-095-130) or CD8a+ T Cell Isolation Kit (mouse, Cat. 130-104-075), respectively. Isolated T cells were resuspended in cRPMI (RPMI-1640 + 10% FBS + 2mM L-Glutamine + 100 U Pen/Strep (Fisher) + 49 nM β-mercaptoethanol (Sigma)) to a final concentration of 1 × 10^6^ cells/ml. Medium for in vivo experiments was supplemented with 2ng/ml IL-2 + 2.5 ng/ml IL-7 + 50 ng/ml IL-15 + 1 μg/ml anti-CD28. Medium for in vitro experiments was supplemented with 2 ng/ml IL-2 + 1 μg/ml anti-CD28. Cells were cultured on plates pretreated with 5 μg/ml anti-CD3. Cytokines and antibodies mentioned above were purchased from BioLegend.

### Human *TRAC* locus knock-in CAR-T cell generation

The CAR-T cells were generated as previously described ^24^. Briefly, 2 million human T cells were mixed with 10 ug Cpf1 mRNA in 110μLof buffer R (Neon Transfection System Kits) and electric-shocked with a 100μLtip under program 24 (1,600 V, 10 ms, and 3 pulses). After electroporation, the cells were immediately transferred into 1 ml of prewarmed X-VIVO media (without antibiotics) immediately. AAV vectors containing gRNA and the CAR construct were added to the T cells 4 hours post-electroporation.

### Human CAR-T cell generated by lentivirus transduction

For human T cells, 10μLof concentrated anti-HER2 CAR lentivirus was added to 2 million human T cells with polybrene (1:1000) in 2 mL X-VIVO media. Cells were spun at 900 g for 90 min before placing back to the incubator. 24 hours later, 1 μg/mL puromycin (Gibco) were added to the media. Puromycin selection was continued until uninfected control cells were no longer viable as determined by flow cytometry.

### xCelligence Real-Time Cell Analysis (RTCA) for co-culture assay

The RTCA E-plates (Agilent) were prepared with 50 µL medium for blanking. 2 × 10^4^ -A498, ACHN, or HT29 cells in additional 50 µL medium were added into each well, and cell index was recorded overnight using the xCELLigence RTCA MP instrument (Agilent) to monitor tumor cells growth. The next day, 100 µL CAR-T cells, ADC, or free drug, with the indicated number were added into each well of the plate. Cell index values were measured every 15 min to evaluate tumor killing. Cell index was normalized at the time point of CAR-T cells addition as normalized cell index.

### Metabolic Labeling of Ac4ManNAz and Carboxyrhodamine 110 in CAR-T cells

CA9 CAR-T cells were seeded on a 96-well plate at a density of 3 × 10^5^ cells/well in 200μLof medium containing the specified concentration series of Ac_4_ManNAz (Vector Labs, Cat. CCT-1084-5). After incubation for 2 days, cells were washed twice with Dulbecco’s phosphate-buffered saline (DPBS, pH 7.4) and subsequently incubated with 200 μl medium containing Carboxyrhodamine-110-PEG4-DBCO (Broadpharm, Cat. BP-22454) at the indicated concentration for 1 h. Following incubation, the cells were washed with DPBS and prepared for subsequent experiments.

### Confocal Microscopy

CAR-T cells metabolically Labeled with Ac_4_ManNAz and Carboxyrhodamine 110-PEG4-DBCO were stained with Hoechst 33342 (ThermoFisher, Cat. H3570) and seeded onto an 8-well chambered coverslip (Ibidi, Cat. 80826). Confocal images were obtained and analyzed using a Leica SP8 Laser Confocal Microscope.

### Time-lapse confocal imaging

CAR-T cells were labeled with Vybrant DiD cell-labeling solution to enable their visualization. Subsequently, these CAR-T cells were conjugated with DBCO-MMAE-Cy7 to facilitate drug dynamic trafficking imaging. Cancer cells were labeled with Vybrant DiO dye, following the manufacturer’s precise protocol.

For the experimental setup, cancer cells were seeded at a density of 20,000 cells per well on an Ibidi 8-well μ-Slide. This seeding was performed one day before imaging to ensure proper cell attachment and growth. The CAR-T-D-C cells were then formed and carefully added on top of the pre-seeded cancer cells. Time-lapse imaging was conducted using a Leica Stellaris confocal microscope. Images were captured at 20X magnification, with acquisition intervals set to every 8 minutes. This continuous imaging was conducted for a total duration of 28 hours.

### Bulk RNA seq sample preparation

CAR-T cells were seeded at 3 × 10^5 cells per well in a 96-well plate and treated with DMSO, Ac4ManNAz, or MMAE as indicated. Following treatment, cells were harvested and total RNA was extracted using TRIzol reagent (Thermo Fisher). RNA-seq libraries were prepared according to the manufacturer’s instructions using the NEBNext rRNA Depletion Kit v2 and the NEBNext Ultra II Directional RNA Library Prep Kit for Illumina.

### Bulk mRNA-seq analysis

Raw sequencing data in FASTQ format underwent initial quality assessment using FastQC (v0.12.1) to evaluate read integrity and detect potential sequencing artifacts. High-quality reads were aligned to the GRCh38 (v45) reference genome using the STAR aligner (v2.7.11a). Gene-level quantification was performed using Gencode v45 annotations, ensuring accurate mapping to known transcript features. Post-alignment, gene count matrices were generated and merged across all samples using a custom R script. Genes with fewer than 20 cumulative counts across all experimental conditions were classified as non-expressed and excluded from downstream analyses to reduce noise and improve statistical power. The filtered count matrix was normalized using DESeq2’s variance-stabilizing transformation, followed by differential expression analysis. Quality control and sample clustering were assessed via Principal Component Analysis (PCA) and unsupervised hierarchical clustering heatmaps, enabling identification of batch effects and biological variability. Differentially expressed genes (DEGs) were defined by a threshold of |Log2FoldChange| > 0.5 and adjusted p-value < 0.05, ensuring robust statistical significance. Functional enrichment of DEGs was performed using Gene Set Enrichment Analysis (GSEA) and the ClusterProfiler R package, leveraging curated pathway databases to identify biologically relevant processes.

### Generation of MMAE CAR-T-D-C

CA9 and HER2 CAR-T cells were seeded on 96-well plates at a density of 3 × 10^5^ cells/well in 200μLof medium containing 20 μM Ac_4_ManNAz (Vector Labs, Cat. CCT-1084-5) and incubated for two days. Following incubation, cells were washed twice with DPBS (pH 7.4) and then treated with 200 μL of medium containing 20 μM DBCO-PEG4-Val-Cit-PAB-MMAE (Broadpharm, Cat. BP-22454) for 1 hour. Following this treatment, the drug was thoroughly washed away, and the resulting MMAE CAR-T-D-C cells were prepared for subsequent experiments.

### Assessment of cell viability

Unless indicated otherwise, 5 × 10^3^ cancer cells or 5 × 10^4^ T cells were seeded in 96-well plates and treated with DBCO-PEG4-Val-Cit-PAB-MMAE (Broadpharm, Cat. BP-22454, DBCO-MMAE for short) or ADC (MCE HY-164992, Trastuzumab vedotin) at the concentration as specified in the figures and legends. DBCO-MMAE was added to the cells the day before adding alamarBlue cell viability reagent (ThermoFisher) to a final concentration of 50 μM. Following 6–8 hours of incubation, cell viability was assessed by recording fluorescence at 540 nm excitation and 590 nm emission using an EnVision 2104 Multilabel Plate Reader (PerkinElmer). Data from at least four replicate wells were averaged for each condition, with cell viability expressed as a percentage relative to the untreated or vehicle-treated control (mean ± SD).

### Assessment of Val-Cit Linker Cleavage in Cell Supernatants Using LC/MS

Cells were cultured for 24 h and the supernatant was harvested to assess val-cit linker cleavage capability. A total of 100μLsupernatant was incubated with 2uM DBCO-PEG4-Val-Cit-PAB-MMAE at 37 °C for 3 h. Following incubation, 50μLof the cell supernatants was mixed with an equal volume of methanol (Thermo Fisher Scientific) to quench the reaction. The sample mixtures were centrifuged at 13,000 rpm for 10 minutes at 4 °C using an Eppendorf Centrifuge 5430R. The resulting supernatants (5 μL) were analyzed through High-Performance Liquid Chromatography coupled with High-Resolution Electrospray Ionization Mass Spectrometry (HPLC-HR-ESI-MS) to perform targeted metabolomics analysis. MS data were obtained using an Agilent iFunnel 6550 quadrupole time-of-flight mass spectrometer (Q-TOF-MS) equipped with an ESI source, coupled to an Agilent 1290 Infinity HPLC system and a Phenomenex Luna C18(2) column (100 Å, 10 μm, 10.0 × 250 mm). A gradient elution containing water (Millipore Sigma) with 0.1% formic acid (Thermo Fisher Scientific) and acetonitrile (Millipore Sigma) with 0.1% formic acid was used at a flow rate of 0.7 mL/min: 0–10 minutes, 10% to 100% acetonitrile with 0.1% formic acid. Extracted ion chromatograms (EICs) with a ±10 ppm window for monomethyl auristatin E (MMAE, [M+H]+ ion at m/z 718.5113, retention time= 6.7 min) were analyzed. The peak areas (area under the curve, AUC) were integrated. MassHunter Qualitative Analysis (v.B.06.00) was used to analyze the MS datasets.

### Synthesis of DBCO-MMAE-Cy7

DBCO-PEG4-val-cit-PAB-MMAE (20 mg, 0.012 mmol) was added to a solution of Cyanine7 carboxylic acid (10.4 mg, 0.018 mmol), EDCI (3.44 mg, 0.018 mmol), 4-Dimethylaminopyridine (1mg, 0.008mmol) and Et_3_N (2.4 mg, 0.024 mmol) in DCM (0.5 ml), which was stirred overnight at room temperature. The mixture was concentrated under vacuum and purified by the flash column chromatography using a gradient eluent of 2% - 6% MeOH in DCM. The product was purified again on preparative thin-layer chromatography to get pure DBCO-MMAE-Cy7. High-resolution mass spectra was used for structural characterization.

### High-resolution mass spectrometry

Mass spectrometric measurements were performed on a Shimadzu Scientific Instruments quadrupole time-of-flight (QToF) 9030 LC-MS system. UV data was collected with a Shimadzu Nexera HPLC/UHPLC Photodiode Array Detector SPD M-40 in the range of 190 - 800nm. Samples were held in the autosampler compartment. LC was performed on a Shim-pack Scepter C18-120, 1.9 μm, 2.1x100 mm Column, equilibrated at 4℃ in a column oven. A gradient eluent with Acetonitrile and Water (0.1% FA) in a mode of 10 min was applied. Flow was held constant at 0.4000 mL/min and the composition of the eluent was changed according to the following gradient: 0 to 2 min, held at 95% A, 5% B; 2 to 8 min, change to 5% A, 95% B; 8 to 10 min, held at 5% A, 95% B; 10 to 10.01 min, change to 95% A, 5% B; 10.01 to 12min, held at 95% A, 5% B. The ionization source was run in “ESI” mode, with the electrospray needle held at +4.5kV. Nebulizer Gas was at 2 L/min, Heating Gas Flow at 10 L/min. Mass spectra were recorded in the range of 100 to 2000 m/z in positive ion mode. Measurements and data post-processing were performed with LabSolutions 5.97 Realtime Analysis and PostRun. The LC-MS tests were completed by the Chemical and Biophysical Instrumentation Center at Yale University.

### Mouse CAR-T cell generation

Mouse CAR-T cells were generated by retrovirus transduction. To produce retrovirus, 4 million HEK293 cells were cultured into 15 cm plate in D10 medium. The following day, the medium was replaced 2 h before transfection. A solution containing 16 μg of transfer plasmid and 8 μg of pCL-Eco packaging plasmid was added to 1 mL of DMEM media. In a separate tube, 62.5 μL of LipoD293 was added to 1 mL of DMEM media and vortexed prior to being added to the DNA:DMEM mixture. The transfection mixture was incubated at room temperature for 10 min before being added to HEK293 cells. Retroviral supernatant was collected 48 hours after transfection, cells were spun down at 3000 g for 15 min to remove cell debris, and then the supernatant were filtered through 0.22 μm filter. The retrovirus was then concentrated by diluting autoclaved 40% PEG8000 (m/v) to a final concentration of 8% PEG8000 in the viral supernatant, followed by overnight incubation at 4 °C. The next day, the retroviral mixture was centrifuged at 1500 g for 30 min, the supernatant was removed, and the pegylated viral pellet was resuspended in 1 mL of complete RPMI (RPMI + 10% FBS + 1% L-glutamine + 1% Pen/Strep + 50 μM β-mercaptoethanol). Retroviral aliquots were stored at −80 °C for future use. Mouse T cells were isolated using the Pan T kit following the manufacturer’s instruction and activated with anti-CD3 and CD28 antibody for 24 hours. Then, 10μLof concentrated CA9-CAR mouse retrovirus was added to 2 million activated mouse T cells in 2 mL media. Cells were spun at 900 g for 90 min before placing back to the incubator. CAR expression was tested 4 days later.

### Establishment of CA9 overexpression cell line via lentivirus transduction

To produce lentivirus, 1 million Lenti-X 293T cells (Invitrogen) were plated into a 6-well plate the night before in D10 medium. The following day, the medium was refreshed, and cells were incubated at 37°C for 1–2 hours prior to transfection. For transfection, 1 μg of transfer plasmid, 0.75 μg of psPAX2 packaging plasmid, and 0.5 μg of pMD2.G envelop plasmid were added to 50 μl of DMEM media. In a separate tube, 3 μl of LipoD293 (Signagen) was diluted in 50 μl of DMEM and then quickly added to DNA:DMEM mixture. The transfection mixture was incubated at room temperature for 10-15 min before being added dropwise to the HEK293 cells. The lentiviral supernatant was collected 48 hours post-transfection, centrifuged at 3000 g for 5 min, and filtered through a 0.22 μm filter. Lentivirus was concentrated using Lenti-X Concentrator, resuspended in media, and either used immediately or stored at −80°C until needed. For transduction of Expi293 and Colon26 cells, 10 μL of concentrated CA9-overexpressing lentivirus was added to 2 million cells in 3 mL of medium with polybrene (1:1000). After 24 hours, 5 μg/mL of puromycin (Gibco) was added to the medium, and selection was continued until all uninfected control cells were nonviable, as confirmed by flow cytometry.

### Human kidney cancer cell line mouse model

NSG mice were subcutaneously injected with 4 million A498 or ACHN renal cell cancer cells on day 0. Subsequently, on days 34 and 40 for the A498 cohort, and on days 17 and 38 for the ACHN cohort, 6 million anti-human CA9-CAR-T/CAR-T-D-C cells, or PBS were intravenously injected into the mice at indicated time point. Tumor progression was evaluated by tumor volume measurement by caliper, calculated as the formula: vol = π/6 ∗ length ∗ width ∗ height.

### Human colon cancer cell line mouse model

Subcutaneous model: NSG mice were subcutaneously injected with 2 million HT29 cancer cells on day 0. On day 15, 2 million anti-human CA9 or anti-human HER2-CAR-T cells, with or without conjugation, or PBS were intravenously injected into the mice. Tumor progression was monitored as mentioned above until mice were sacrificed when tumor reached 2000 mm^3^.

Lung metastasis model: NSG mice were intravenously injected with 1 × 10⁶ HT29-GL cells via the tail vein. Mice were randomized to treatment groups once tumors were established. Tumor progression was monitored by bioluminescence imaging. At study endpoints, mice were euthanized and lungs collected for downstream analyses.

### Syngeneic colon cancer mouse model

3 million Colon26 cancer cells overexpressed human CA9 (Colon26-CA9) were subcutaneously inoculated into female BALB/c mice on day 0. When tumors reached 100 mm^3^, 5 million anti-CA9-CAR-T/CAR-T-D-C cells, or PBS were intravenously injected into the mice. Tumor progression was monitored as previously described. Additionally, tumors were harvested on day 7 post-treatment for flow cytometry analysis of tumor-infiltrating lymphocytes.

### Syngeneic melanoma mouse model

2.5 million mouse B16F10-OVA cells were subcutaneously inoculated into female C57BL/6 mice on day 0. On day 6, OT-I T cells were isolated from OT-I mice spleen, then in vitro expand for 5 days. On day 11, 7 million OT-I T cells, with or without conjugation, or PBS were intravenously injected into the mice. Tumor progression was monitored as mentioned above. For tumor-infiltrating immune cells profiling, the tumors were harvested on day 7, followed by flow cytometry analysis of tumor-infiltrating lymphocytes. For survival curve depicting, mice were monitored until tumor reached 2000 mm^3^.

### Syngeneic bilateral tumors models

Two models were used for studying section. For the **Colon26 model**, BABL/c mice were subcutaneously injected with 3 million mouse Colon26-CA9 cells in the right flank and 2 million mouse Colon26 cells in the left flank. When tumors reached 100 mm^3^, 5 million anti-CA9 mouse CAR-T cells, with or without conjugation, or PBS were intravenously injected into the mice. For the **Melanoma model**, C57BL/6 mice were subcutaneously injected with 3 million mouse B16F10-OVA cells in the right flank and 2 million mouse B16F10 cells in the left flank. On day 11, 7 million OT-I T cells, with or without conjugation, or PBS were intravenously injected into the mice. Tumor progression was monitored as mentioned above. Tumors were harvested at the specified endpoints, followed by ELISPOT assays to assess immune responses.

### Spatial Sample Handling and Section Preparation

One million HT29-GL cells were intravenously injected into NSG mice, and tumor progression was monitored using IVIS imaging. On day 21 post-injection, the mice were treated with 5 million CAR-T or CAR-T-D-C cells. The lungs were harvested 24 hours post-treatment for spatial transcriptomics. To collect the lung samples, the harvested lungs bearing HT29 tumors were preserved with Optimal Cutting Temperature (OCT) compound and stored at -80°C for future use. The samples were sectioned at a thickness of 7-10 μm and centrally mounted on Poly-L-Lysine coated 1 x 3” glass slides. Serial tissue sections were collected simultaneously for Digital Barcoding in Tissue (DBiT) and Hematoxylin and Eosin (HE) staining. Sample preparation for transcriptomic sequencing was performed as previously described^43^, with key steps summarized below.

### DNA Barcodes Annealing

DNA oligos (Table S3) were obtained from IDT. Barcodes (100 µM) and ligation linkers (100 µM) were annealed 1:1 in 2× annealing buffer (20 mM Tris-HCl pH 8.0, 100 mM NaCl, 2 mM EDTA) using the program: 95 °C for 5 min, slow cooling to 20 °C at 0.1 °C/s, followed by 12 °C for 3 min. Annealed barcodes were stored at −20 °C until use.

### Permeabilization, In Situ Polyadenylation, and Reverse Transcription

Permeabilization, In Situ Polyadenylation, and Reverse TranscriptionTissue slides were permeabilized with 1% Triton X-100 in DPBS for 20 min at room temperature, then washed with 0.5× DPBS-RI (1× DPBS diluted with nuclease-free water, 0.05 U/µL RNase inhibitor). A PDMS reservoir was attached to cover the ROI. In situ polyadenylation was performed using E. coli Poly(A) Polymerase in 60 µL reaction mix at 37 °C for 30 min, followed by a DPBS wash. Reverse transcription was performed in 60 µL RT mix at room temperature for 30 min, then 42 °C for 90 min, followed by washing with DPBS.

### Spatial Barcoding with Microfluidic Devices

The first PDMS device was aligned over the ROI and secured with an acrylic clamp. Barcode A (25 µM) was ligated using T4 DNA ligase mix, introduced into the channels, and incubated at 37 °C for 30 min. The device was removed, and the slide washed with DPBS. Barcode B was ligated using a perpendicular PDMS device in the same manner. After five washes with nuclease-free water, the microchannel pattern was recorded.

### Tissue Lysis and cDNA Extraction

The ROI was enclosed with a PDMS reservoir and 70 µL lysis mix (1× DPBS, 2× lysis buffer, Proteinase K) was added and incubated at 55 °C for 2 h. An additional 40 µL lysis mix was applied to recover residual cDNA, and lysates were incubated overnight at 55 °C to reverse crosslinks. Lysates were stored at −80 °C until use.

### cDNA Purification, Template Switch, and PCR Amplification

Proteinase K was inactivated with 5 µL PMSF (100 µM). cDNA was purified using Dynabeads MyOne Streptavidin C1 in 2× B&W buffer, washed, and subjected to template-switch reaction at room temperature for 30 min and 42 °C for 90 min. Beads were washed, resuspended in PCR mix, and amplified with an initial program (95 °C 3 min; 5 cycles of 98 °C 20 s, 63 °C 45 s, 72 °C 3 min; 72 °C 3 min). qPCR determined additional cycles, and PCR products were purified with SPRIselect beads (0.8×) and analyzed on TapeStation D5000.

### rRNA Depletion, Library Preparation, and Sequencing

20 ng cDNA underwent three rounds of rRNA and mitochondrial rRNA depletion using the SEQuoia RiboDepletion Kit. Sequencing adapters were ligated using 7 PCR cycles (2× KAPA HiFi, P5/P7 primers), purified (0.8× SPRIselect), quality-checked, and sequenced on Illumina NovaSeq 6000 (paired-end 150 bp).

### Spatial sequence alignment and generation of mRNA expression matrix

To decode sequencing data, Read 2 FASTQ files were processed to extract unique molecular identifiers (UMIs) and spatial barcodes A and B. Read 1, containing cDNA sequences, was trimmed with Cutadapt and aligned to the human reference genome (GRCh38) using STAR (v2.7.11b). Spatial barcodes were demultiplexed with ST_Pipeline (v1.7.6) according to the predefined microfluidic channel coordinates, generating a gene-by-pixel expression matrix for downstream analyses. Matrix entries corresponding to pixels without tissue coverage were excluded.

### Isolation of tumor infiltrating immune cells

Syngeneic Colon26 and B16F10 tumors were established by intravenous transplantation. Tumor-bearing mice were randomly assigned to treatment groups as indicated in the figure legends. Mice were euthanized 7 days post-treatment, and tumors were harvested into ice-cold 2% FBS. Tumors were minced into 1–3 mm fragments using a scalpel and digested with 100 U/mL Collagenase IV for 30–60 min at 37 °C with constant stirring. Tumor suspensions were sequentially filtered twice through 100 µm strainers and once through a 40 µm strainer to remove debris and large aggregates. Red blood cells were lysed with 1 mL ACK Lysis Buffer (Lonza) per spleen for 2–5 min at room temperature, followed by dilution with 10 mL 2% FBS and filtration through a 40 µm strainer. The resulting single-cell suspensions were used for flow cytometry staining.

### Flow cytometry analysis of tumor infiltrated immune cells

All antibodies used for flow cytometry were purchased from BioLegend or eBioscience. Single-cell suspensions of tumors or spleens were prepared using a gentleMACS tissue dissociation system. The following antibody panels were used: Panel 1: Live/death-APC/Cy7, anti-CD45.2-PerCP/Cy5.5, anti-CD3-PE, anti-CD4-PE/Cy7, anti-CD8a-Bv605, anti-PD1-APC, anti-CCR7-FITC and CD103-Bv421; Panel 2: Panel 1: Live/death-APC/Cy7, anti-CD45.2-PerCP/Cy5.5, anti-CD3-PE, anti-CD4-PE/Cy7, anti-CD8a-Bv605, anti-PD1-APC, anti-IFN-γ-BV421; Panel 3: Live/death-APC/Cy7, anti-CD45.2-PerCP/Cy5.5, anti-CD11b-APC, anti-Ly6c-BV605; anti-F4/80-PE, and anti-CD11c-PE/Dazzle594. Unless otherwise specified, antibodies were used at a 1:200 dilution. For intracellular staining, cells were processed using the eBioscience™ Intracellular Fixation & Permeabilization Buffer Set according to the manufacturer’s instructions. Briefly, after surface marker staining, cells were resuspended in 100 μl Fixation/Permeabilization working solution and incubated on ice for 10 min. Cells were washed once with 1× Permeabilization Buffer, centrifuged at 600 g for 5 min, resuspended in 50 μl 1× Permeabilization Buffer containing Fc receptor block, and incubated on ice for 10 min. Intracellular antibodies (diluted 2× in Permeabilization Buffer) were then added in 50 μl volume, and cells were incubated on ice for 30 min. Following staining, cells were washed twice with staining buffer and analyzed or sorted on a BD FACSAria. Data were analyzed using FlowJo software. Identification of monocyte, neutrophil, dendritic cell, and macrophage populations was based on a previously described gating strategy, with modifications^44^ .

### IFN-gamma ELISPOT assay

Mouse spleens were harvested at the specified time points from the indicated mouse strains and placed in ice-cold washing buffer (2% FBS in PBS). The spleens were then mashed through a 100 μm filter in washing buffer. Red blood cells were lysed using ACK lysis buffer, incubated for 2 min at room temperature, and subsequently washed with washing buffer. The cell suspensions were filtered through a 40 μm filter and resuspended in cRPMI medium. Following cell counting, 0.1 million splenocytes were seeded per well in 100 µL of cRPMI. An additional 0.1 million tumor cells were added to each well for co-culture over a 48-hour period. Imaging and analysis were conducted using an ImmunoSpot S6 Entry M2 plate reader.

### Protein structure in schematic diagram

Schematic diagram comparing CA9 and HER2 was created with Biorender. Protein structures were obtained from Protein Data Bank (PDB). We use 6FE2 to represent the structure of CA9 and 5O4G for the structure of HER2 and its binding Fab.

### TCGA data retrieve and analysis

Curated TCGA dataset was obtained through Broad GDAC Firehose, using API FirebrowseR.

Specifically, mRNA expression data regarding the gene of interest (CA9 and CTSB) was retrieved. Only patient samples with valid expression_log2 and both tumor & tumor adjacent normal tissue data were kept. The dataset was further filter out cohort with less than 10 patient samples. Expression_log2 data was visualized using ggplot2.

### Illustrations

Illustration of schematics were performed using Adobe Illustrator and Biorender (https://www.biorender.com).

### Standard statistical analysis

Data analysis was performed using GraphPad Prism version 10 and R Studio. Detailed statistical methods are provided in figure legends and/or supplementary Excel tables of Source data. Different levels of statistical significance were assessed based on specific *p*-values and type I error cutoffs (* 0.05, ** 0.01, *** 0.001 and **** 0.0001).

**Source data and statistics** are provided in an excel file.

## Data and resource availability

All generated data and analysis information/results for this this study are included in this article and its supplementary information files. Raw sequencing data are being deposited to the Gene Expression Omnibus (GEO). Original cell lines are available at the commercial sources listed in methods and/or reporting summary. Other relevant information or data are available from the lead corresponding author (SC) upon reasonable request.

## Supplemental Figure Legends

**Figure S1. Generation of human anti-CA9 CAR-T binder using mRNA-LNP immunization and single-sell BCR sequencing.**

**(a)** Analysis from The Cancer Genome Atlas (TCGA) showing the expression level of *CA9* mRNA across different cancer cell types.

**(b)** Illustration depicting the structural properties of CA9 that prevent endocytosis, rendering it less suitable as a target for antibody-drug conjugate (ADC) due to the lack of internalization.

**(c)** Histogram showing A498 cancer cell lysis following co-culture with T cells or G250-CAR-T cells. Cytolysis (%) was calculated from the reduction in luciferase signal relative to untreated cancer cells.

**(d)** Histogram showing the radius distribution of CA9 LNP-mRNA.

**(e)** ELISA demonstrating the reactivity of serum antibody in mice plasma sample collected on day14 (prime) and day 21 (booster) with the human CA9 target antigen. The samples were diluted with PBS as indicated. Data are shown as mean ± SD plus individual data points in dot plots (n = 2). Statistical analysis was performed using two-way ANOVA. Statistical significance is denoted as follows: ∗∗∗∗p < 0.0001.

**(f)** Distribution of BCR variable region V, D, and J gene segment usage. Sequencing analysis of 1,407 CA9-specific B cells yielded 627 paired antibody sequences.

**(g)** Frequency of BCR gene usage of the sequenced memory B cells.

**Figure S2. Binding of anti-CA9 antibodies to CA9 antigen and the manufacturing process of anti-CA9 CAR-T cells.**

**(a)** Flow cytometry showing the binding ability of the top anti-CA9 clones in human IgG format with Expi293F cells overexpressing CA9. A commercial anti-human CA9 antibody (CA9-PE) was used to verify CA9 overexpression. Sequence from a prior binder (clone G250) was used as a positive control.

**(b)** Schematics showing the anti-human CA9 CAR-T cell production procedure.

**(c)** CAR expression on the cell surface was confirmed by Flag detection via flow cytometry. The scFvs used for CAR-T were selected based on their binding efficacy as shown in Figure 1c.

**Figure S3. Determination of the optimal dose for glycolabeling and assessment of labeling efficiency across multiple cell types.**

**(a)** Histogram showing the fluorescence intensity measured with flow cytometry in the AlexaFluor 488 channel. Human CAR-T cells were treated with varying concentrations of Ac4ManNAz (ranges from 0 to 80 μM) and DBCO-PEG4-Carboxyrhodamine 110 (ranges from 5 to 50 μM).

**(b)** Bar plot showing mean fluorescence intensity (MFI) of human CAR-T cells treated with different concentrations of Ac4ManNAz and DBCO-PEG4-Carboxyrhodamine 110.

**(c)** Heat maps showing the MFI and Carboxyrhodamine 110 percentage for each concentration combination of Ac4ManNAz and DBCO-PEG4-Carboxyrhodamine 110 to determine the saturation level.

**(d)** Statistics analysis of the fluorescence intensity in human CAR-T cells treated with 20μMAc4ManNAz and indicated concentration of DBCO-PEG4-Carboxyrhodamine 110. Data are presented as mean ± SD. Statistical significance was assessed using two-way ANOVA. Statistical significance labels: ∗p < 0.05; ∗∗p < 0.01; ∗∗∗p < 0.001; ∗∗∗∗p < 0.0001. MFI: median fluorescence intensity.

**(e)** Volcano plot depicting differential mRNA profiles of Ac₄ManNAz–treated versus DMSO control T cells from RNA-seq data.

**(f)** Flow cytometry gating strategy for PBMCs labeled with Ac₄ManNAz–Carboxyrhodamine 110.

**(g)** Histogram showing fluorescence intensity of various cell types after glycolabeling. Human PBMC were treated with 20 μM Ac4ManNAz and 20 μM DBCO-PEG4-Carboxyrhodamine 110.

**Figure S4. Assessment of cell type–specific sensitivity to DBCO-MMAE treatment.**

**(a)** Schematic diagram depicting the in vitro process used to assess DBCO-PEG4-Val-Cit-PAB-MMAE (DBCO-MMAE) sensitivity

**(b)** The viability of four different cancer cell lines and human T cells following treatment with DBCO-MMAE for 24h.

**(c)** Flow cytometry analysis of cytokine release from CAR-T cells treated with free MMAE.

**(d-f)** Volcano plots showing transcriptomic profiles of CAR-T cells treated with free MMAE at 6 h **(d)** and 24 h **(e)**, with Δ-expression per hour calculated to reflect temporal changes **(f)**.

**Figure S5. Verification and quantification of CTSB mediated MMAE release.**

**(a-b)** LC-MS analysis confirming val-cit linker cleavage and MMAE release from the CAR-T-D-C module. **(a)** Intensity of the released MMAE product. **(b)** Area under the curve (AUC) quantification for MMAE release. CAR-T-D-C was incubated for 3 h with supernatants from cultured A498 (A498+CAR-T-D-C), ACHN (ACHN+CAR-T-D-C), EMEM+CAR-T-D-C, and EMEM control prior to LC-MS analysis.

**(c)** Standard calibration curve for MMAE quantification by LC–MS.

**(d)** Diagram of the MMAE conjugation process on CAR-T cells, followed by co-culture assays with tumor cells. Cell killing analysis was performed using the RTCA assay.

**(e-f)** Real-Time Cell Analysis (RTCA) killing assays were employed to assess CAR-T-D-C killing ability against A498 (e) and ACHN (f) cells with or without CTSB treatment, as indicated by normalized cell index values. Vehicle, DMSO vehicle control. E: T, effector cell to target cell ratio.

**(g)** Cytokine release from CAR-T cells following co-culture with A498 cells. Statistical comparisons were made with one-way ANOVA. Statistical significance is denoted as follows: ∗p < 0.05; ∗∗∗p < 0.001; ∗∗∗∗p < 0.0001.

**Figure S6. Characterization of DBCO-MMAE-Cy7 and its targeted delivery to cancer cells.**

**(a)** Chemical structure of DBCO-MMAE-Cy7. The fluorescent Cy7 dye is highlighted in red.

**(b)** UV chromatogram of DBCO-MMAE-Cy7.

**(c)** Mass spectrum of DBCO-MMAE-Cy7. HRMS (ESI) m/z for C125H171N14O20+, calculated 1094.63945 [M++H+], found 1094.6515.

**(d)** Time-lapse confocal imaging showing that cleavage of DBCO-MMAE-Cy7 begins after ∼2 h of co-culture, with the released compound preferentially taken up by cancer cells. DBCO-MMAE-Cy7 acts synergistically with CAR-T cells to kill cancer cells without affecting CAR-T viability. White arrows indicate released DBCO-MMAE-Cy7. Colors: blue, cancer cells; green, CAR-T cells; magenta, DBCO-MMAE-Cy7.

**Figure S7. Expanding the CAR-T-D-C strategy to target additional surface antigens.**

**(a)** Flow cytometry gating strategy for in vivo CAR-T cells, assessing surface marker expression and cytokine release.

**(b-c)** Body weight of mice bearing A498 **(b)** and ACHN **(c)** tumors measured overtime following treatments with PBS vehicle control, CAR-T and CAR-T-D-C.

**Figure S8. Expanding the CAR-T-D-C strategy to target additional surface antigens.**

**(a)** Flow cytometry analysis showing the surface expression of the HER2 antigen on the HT29 cell line

**(b)** Illustration depicting the differential internalization behavior: antibody binding to CA9 does not lead to internalization, whereas antibody binding to HER2 induces internalization.

**(c)** Survival curve of the HT29-GL lung metastasis model.

**(d)** Killing curves of trastuzumab vedotin in HT29 cells were generated at multiple concentrations and timepoints as indicated in the figure, with viability readouts obtained at 24, 48, and 72 hours.

**(e)** Real-Time Cell Analysis (RTCA) killing assays were performed to evaluate the cytotoxic activity of CAR-T-D-C killing ability against HT29 cells in comparison with multiple treatment conditions, including anti-HER2 CAR-T alone (E:T= 1:1, 1:2, 1:5, respectively), MMAE alone (0.25, 0.125, and 0.05 µM, respectively), anti-Her2 CAR-T+MMAE, anti-HER2 ADC alone (trastuzumab vedotin, 0.07, 0.035, and 0.014 µM, respectively), ADC+CAR-T, or DMSO vehicle control. E:T: effector to target ratio. Statistical analysis was performed using two-way ANOVA. Statistical significance is denoted as follows: ∗∗p < 0.01; ∗∗∗p < 0.001; ∗∗∗∗p < 0.0001.

**Figure S9. Spatial transcriptomic for CAR-T cell distribution and function in vivo.**

**(a)** IVIS imaging of tumor progression of the HT29-GL lung metastasis model.

**(b)** Gene expression heatmap of distinct transcriptional clusters of the spatial transcriptomics.

**Figure S10. Spatial transcriptomic mapping of CAR-T cell distribution and function in vivo.**

**(a-b)** Spatial distribution of distinct transcriptional clusters of CAR-T **(a)** and CAR-T-D-C **(b)** groups.

**Figure S11. Flow cytometry gating strategy for tumor-infiltrating immune cells.**

## Notes

### Competing Interest Statement

The authors have declared no competing interest.

